# Predicting mammalian hosts in which novel coronaviruses can be generated

**DOI:** 10.1101/2020.06.15.151845

**Authors:** Maya Wardeh, Matthew Baylis, Marcus S.C. Blagrove

## Abstract

Novel pathogenic coronaviruses – including SARS-CoV and SARS-CoV-2 – arise by homologous recombination in a host cell^1,2^. This process requires a single host to be infected with more than one type of coronavirus, which recombine to form novel strains of virus with unique combinations of genetic material. Identifying possible sources of novel coronaviruses requires identifying hosts (termed recombination hosts) of more than one coronavirus type, in which recombination might occur. However, the majority of coronavirus-host interactions remain unknown, and therefore the vast majority of recombination hosts for coronaviruses cannot be identified. Here we show that there are 11.5-fold more coronavirus-host associations, and over 30-fold more potential SARS-CoV-2 recombination hosts, than have been observed to date. We show there are over 40-fold more host species with four or more different subgenera of coronaviruses. This underestimation of both number and novel coronavirus generation in wild and domesticated animals. Our results list specific high-risk hosts in which our model predicts homologous recombination could occur, our model identifies both wild and domesticated mammals including known important and understudied species. We recommend these species for coronavirus surveillance, as well as enforced separation in livestock markets and agriculture.

## INTRODUCTION

The generation and emergence of three novel respiratory coronaviruses from mammalian reservoirs into human populations in the last 20 years, including one which has achieved pandemic status, suggests that one of the most pressing current research questions is: In which reservoirs could the next novel coronaviruses be generated and emerge from in future? Armed with this knowledge, we may be able to reduce the chance of emergence into human populations or develop potential mitigations in advance, such as by the strict monitoring and enforced separation of the hosts identified here, in live animal markets, farms, and other close-quarters environments.

*Coronaviridae* are a family of positive sense RNA viruses which can cause an array of diseases. In humans, these range from mild cold-like illnesses to the lethal respiratory tract infections mentioned above. Seven coronaviruses are known to infect humans^3^, SARS-CoV, MERS-CoV and SARS-CoV-2 causing severe disease, while HKU1, NL63, OC43 and 229E tend towards milder symptoms in most patients^4^.

Coronaviruses undergo frequent host-shifting events between non-human animal species, or non-human animals and humans^5–7^, a process that may involve changes to the cells or tissues that the viruses infect (virus tropism). Such shifts have resulted in new animal diseases (such as bovine coronavirus (BCoV) disease^8^ and canine coronavirus (CCoV) disease^9^), and human diseases (such as OC43^10^ and 229E^11^). The aetiological agent of COVID-19, SARS-CoV-2, is thought to have originated in bats^12^ and shifted to humans via an intermediate reservoir host, likely a species of pangolin^13^.

Comparison of the genetic sequences of bat and human coronaviruses has revealed five potentially important genetic regions involved in host specificity and shifting, with the Spike receptor binding domain believed to be the most important^1,5^. Homologous recombination is a natural process which brings together new combinations of genetic material, and hence new viral strains, from two similar non-identical parent strains of virus. This recombination occurs when different strains co-infect an individual animal, with sequences from each parent strain in the genetic make-up of progeny virus. Homologous recombination has previously been demonstrated in many important viruses such as human immunodeficiency virus^14^, classical swine fever virus^15^, and throughout the Coronaviridae^1,2^, including homologous recombination in Spike being implicated in the generation of SARS-CoV-2^2^. As well as instigating host-shifting, homologous recombination in other regions of the virus genome could also introduce novel phenotypes into coronavirus strains already infectious to humans. There are at least seven potential regions for homologous recombination in the replicase and Spike regions of the SARS-CoV genome alone, with possible recombination partner viruses from a range of other mammalian and human coronaviruses^16^. Recombination events between two compatible partner strains in a shared host could thus lead to future novel coronaviruses, either by enabling pre-existing mammalian strains to infect humans, or by adding new phenotypes arising from different alleles to pre existing human-affecting strains.

The most fundamental requirement for homologous recombination to take place is the co-infection of a single host with multiple coronaviruses. However, our understanding of which hosts are permissive to which coronaviruses, the pre-requisite to identifying which hosts are potential sites for this recombination (henceforth termed ‘*recombination hosts*’), remains extremely limited. Here, we utilise a similarity-based analytical pipeline to address this significant knowledge gap. Our approach predicts associations between coronaviruses and their potential mammalian hosts by integrating three perspectives or points of view encompassing: 1) genomic features depicting different aspects of coronaviruses (e.g. secondary structure, codon usage bias) extracted from complete genomes (sequences = 3,271, virus strains = 411); 2) ecological, phylogenetic and geospatial traits of potential mammalian hosts (n=876); and 3) characteristics of the network that describes the linkage of coronaviruses to their observed hosts, which expresses our current knowledge of sharing of coronaviruses between various hosts and host groups.

Topological features of ecological networks have been successfully utilised to enhance our understanding of pathogen sharing^17,18^, disease emergence and spill-over events^19^, and as means to predict missing links in a variety of host-pathogen networks^20–22^. Here we capture this topology, and relations between coronaviruses and hosts in our network, by means of node (coronaviruses and hosts) embeddings using DeepWalk^23^ – a deep learning method that has been successfully used to predict drug-target^24^, and IncRNA-disease associations^25^.

Our analytical pipeline transforms the above features into similarities (between viruses, and between hosts) and uses them to give virus–mammal associations scores of how likely they are to occur. Our framework then ensembles its constituent learners to produce testable predictions of mammalian hosts of multiple coronaviruses.

In this study we address the following three questions: 1) Which species may be unidentified mammalian reservoirs of coronaviruses? 2) What are the most probable mammalian host species in which coronavirus homologous recombination could occur? And 3) Which coronaviruses are most likely to co-infect a single host, and thus act as sources for future novel viruses?

## RESULTS

Our model to predict unobserved associations between coronaviruses and their mammalian hosts indicated a total of 126 (ensemble mean probability cut-off>0.5, when subtracting/adding SD from the mean the number of predicted hosts is 85/169. For simplicity, we report SD hereafter as −/+ from predicted values at reported probability cut-offs, here: SD=−41/+43) non-human mammalian species in which SARS-CoV-2 could be found. The breakdown of these hosts by order was as follows (values in brackets represent SD from ensemble mean): Carnivora: 37 (0/0); Rodentia: 32 (−9/3); Chiroptera: 25 (−19/38); Artiodactyla: 20 (−8/2); Eulipotyphla: 5 (−4/0); Primates: 4 (0/0); Lagomorpha: 2 (−1/0) and Pholidota: 1 (0/0). Figure 1 illustrates these predicted hosts, the probability of their association with SARS-COV2, as well as numbers of known and unobserved (predicted) coronaviruses that could be found in each potential reservoir of SARS-CoV-2 (Supplementary results table SR1 lists full predictions).

**Figure 1.**
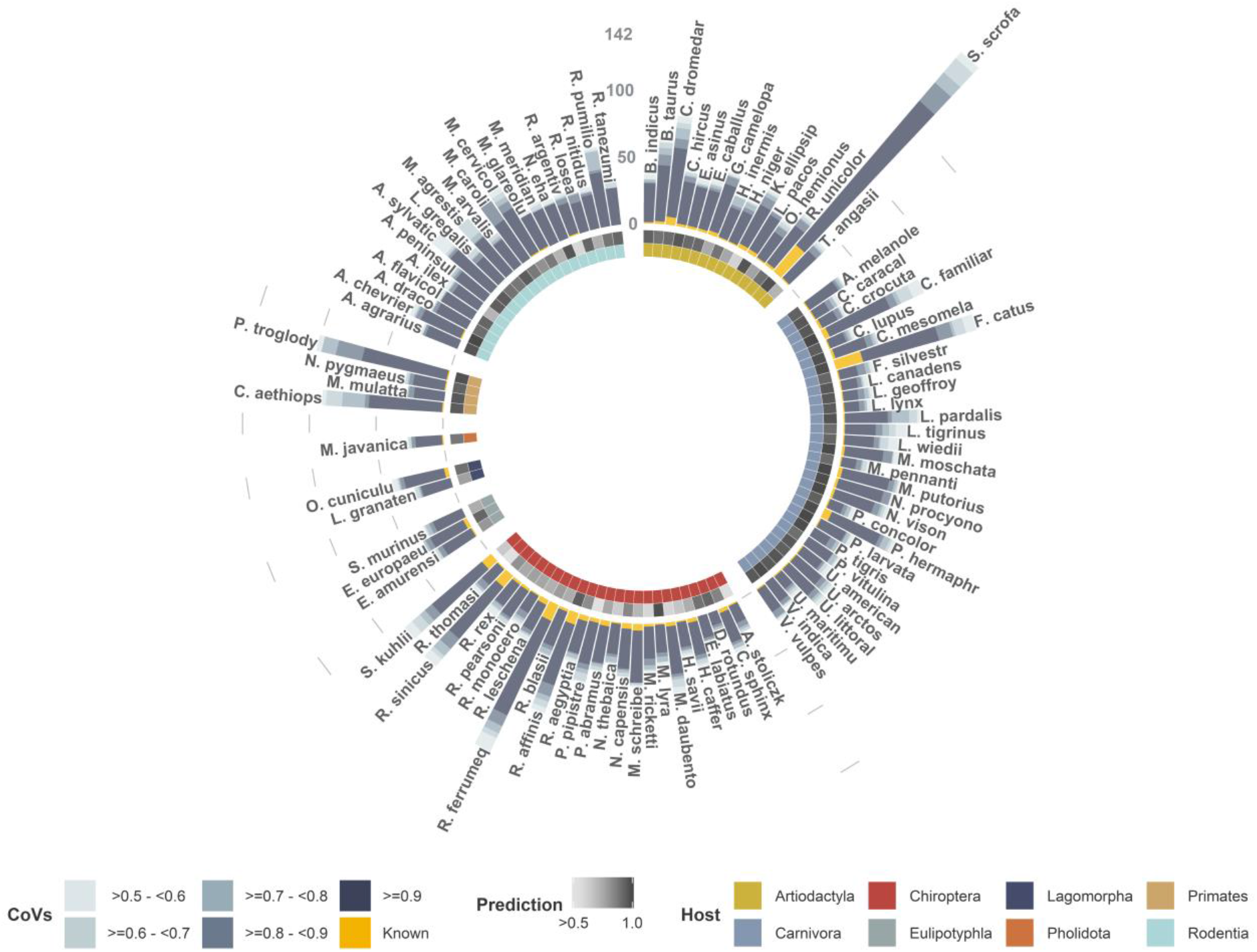
Model predictions for potential hosts of SARS-Cov-2 – excluding humans and lab rodents. Predicted hosts are grouped by order (inner circle). Middle circle presents probability of association between host and SARS-CoV-2 (>0.5 light grey to 1 dark grey). Yellow bars represent number of coronaviruses (species or strains) observed to be found in each host. Blue stacked bars represent other coronaviruses predicted to be found in each host by our model. Predicted coronaviruses per host are grouped by prediction probability into five categories (from inside to outside): >0.9, 0.9-0.9, 0.8-0.7, 0.7-0.6, and 0.6-05.

Overall, our model predicted 4,438 (mean, SD=−1,903/+2,256, cut-off=0.5) previously unobserved associations that potentially exist between 300 (SD=0/+3) mammals and 201 coronaviruses (species or strains, SD=−60/+13). Our model predicts there are 115 (0/+3) mammal species with no previously observed associations with the 411 input viruses.

On average, each coronavirus (species or strain, complete-genome available, N=411) is predicted to have 12.56 mammalian hosts (ensemble mean, SD=−4.92/+5.83, cut-off =0.5). Similarly, each mammalian species (N=876, known hosts =185, predicted hosts= 300, SD=−0/+3) is host to, on average, 5.55 coronaviruses (SD=−2.17/+2.58) – Supplementary tables SR2 and SR3 provide results for each coronavirus and mammalian host, respectively.

Figure 2 presents 50 potential mammalian recombination hosts of coronaviruses. Our model predicts 231 (SD=−115 /+58) mammalian species that could host 10 or more of the 411 coronavirus species or strains for which complete genome sequences were available. The breakdown of these hosts by order was as follows: Chiroptera = 129 (−87/+53); Carnivora: 35 (−22/+2); Rodentia = 33 (−4/+3); Artiodactyla = 22 (−2/+0); Eulipotyphla = 5 (−0/+0); Primates = 4 (−0/+0); Lagomorpha = 2 (−0/+0); and Pholidota =1(−0/+0).

**Figure 2.**
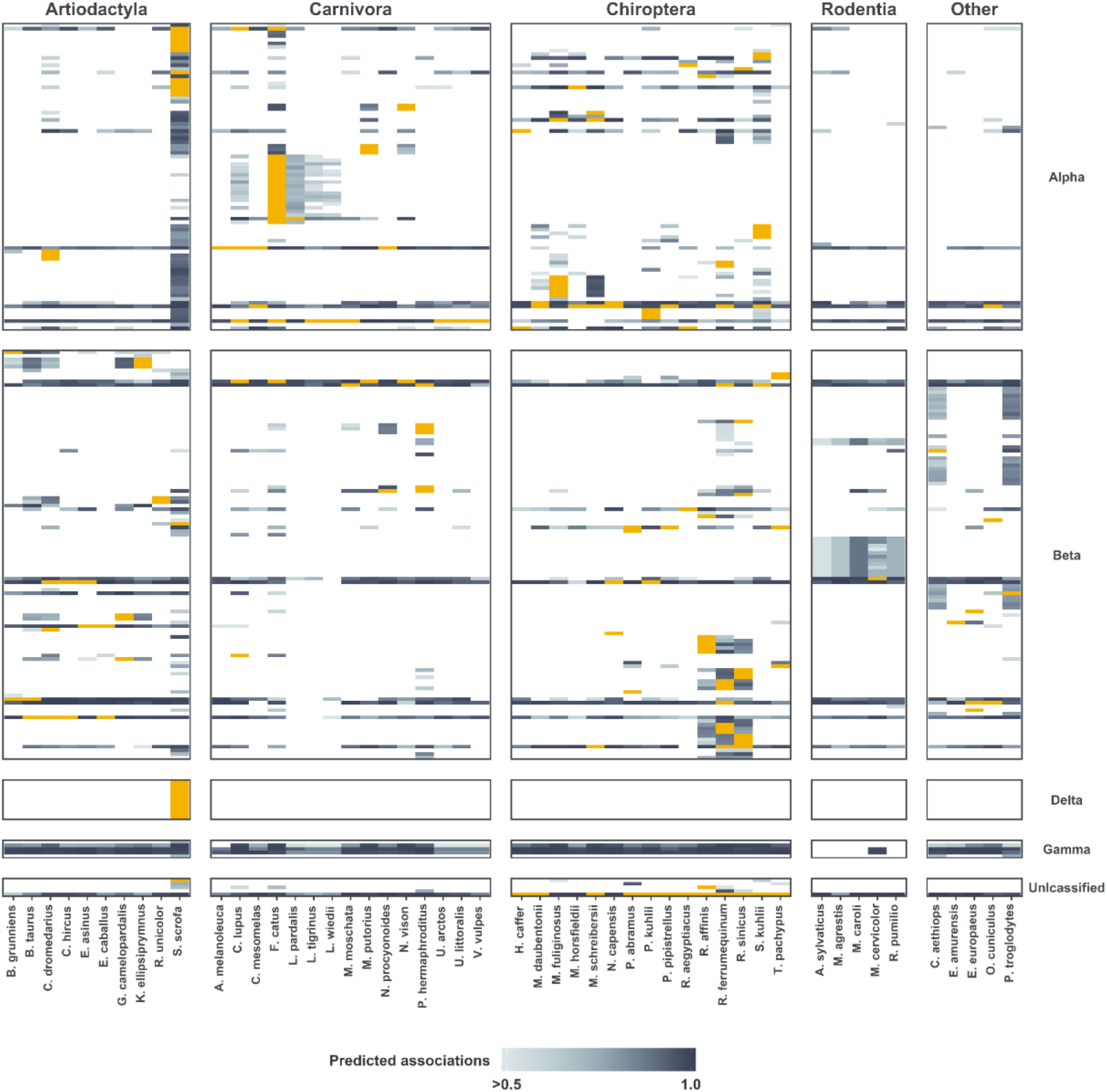
Observed and predicted mammalian hosts for coronavirses. Columns present mammalian hosts in four categories: Artiodactyla (top 10 hosts by number of predicted coronaviruses that could be found in each host); Carnivora (top 15 hosts), Chiroptera (top 15 hosts, each predicted to host 50 or more coronavirus species or strain), and others (top 5). Rows present viruses ordered into five taxonomic groups: Alphacoronaviruses, Betacoronaviruses, Deltacoronaviruses, Gammacoronaviruses and unclassified coronaviruses. Yellow cells represent observed associations between the host and the coronavirus. Blue cells present predicted associations (predicted probability ranging from >0.5 (light blue) to 1 (dark blue)). White cells represent no association between host and virus (beneath cut-off probability of 0.5).

The addition of predicted associations increased the diversity (mean phylogenetic distance) of mammalian hosts per coronavirus, as well as the diversity (mean genetic distance) of coronaviruses per mammalian host (table 1 lists these changes at 0.95, 0.75 and 0.50 probability cut-offs).

**Table 1.**
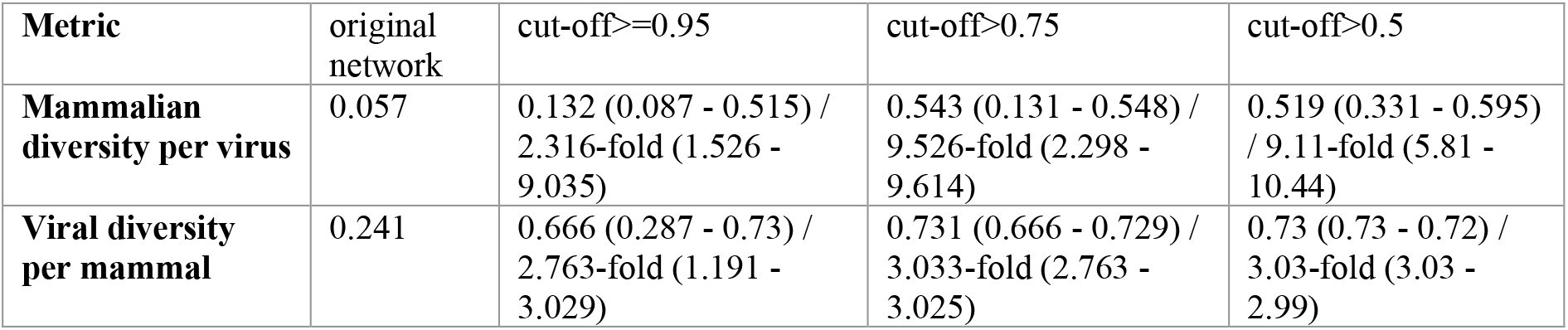
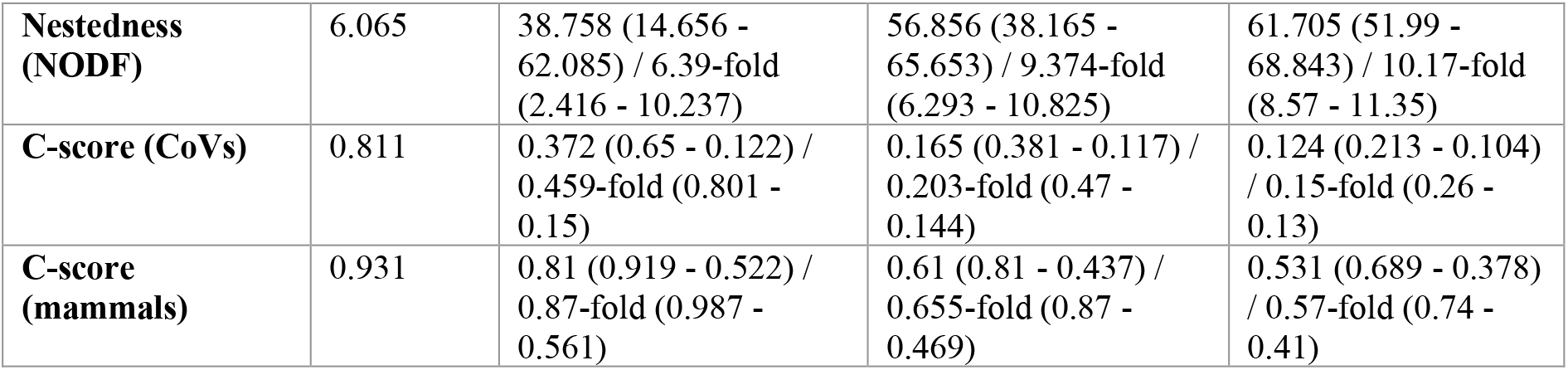
bipartite network metrics calculated for original network, and predicted networks at three probability cut-offs: >0.5, >0.75 and ≥0.95.

**Table 1.**
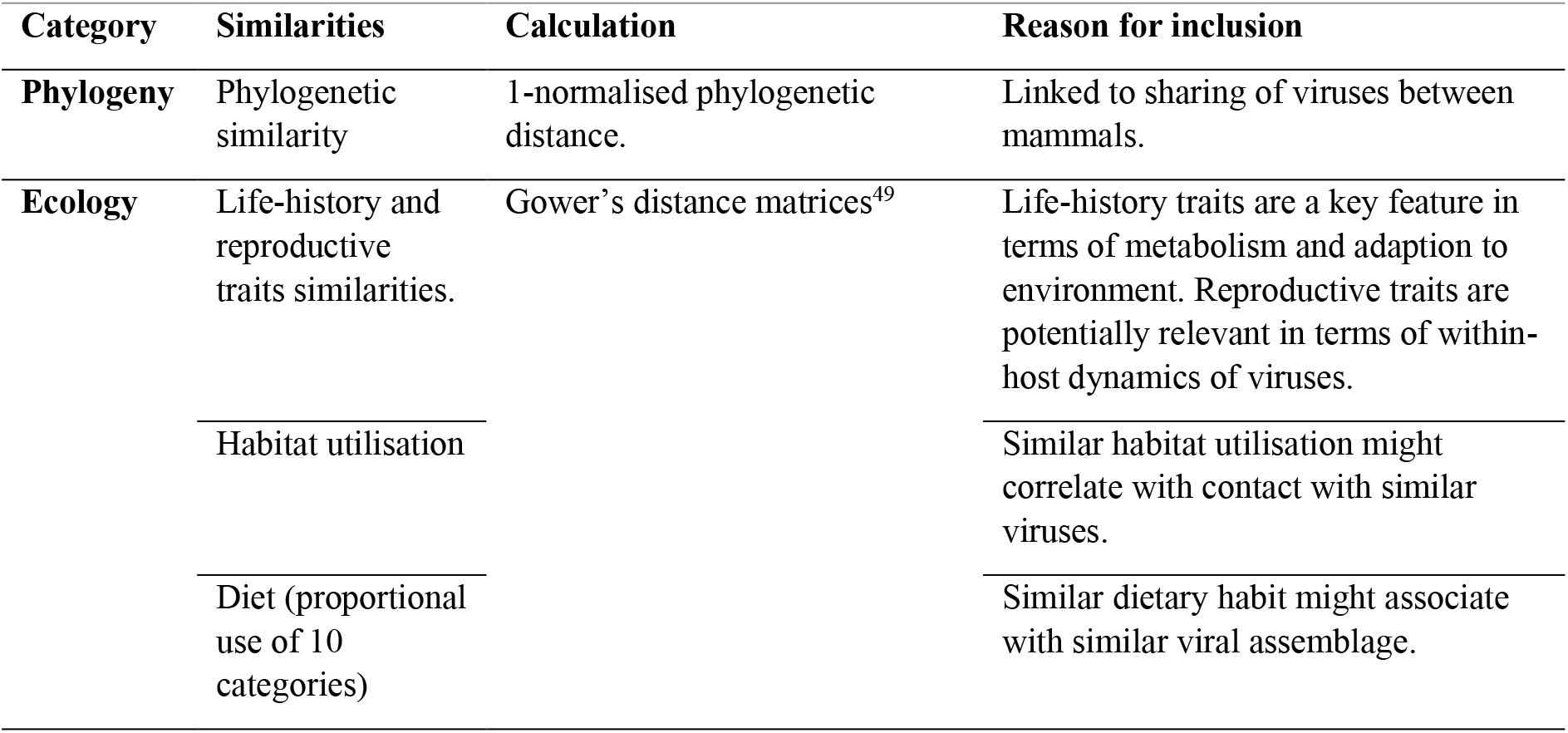
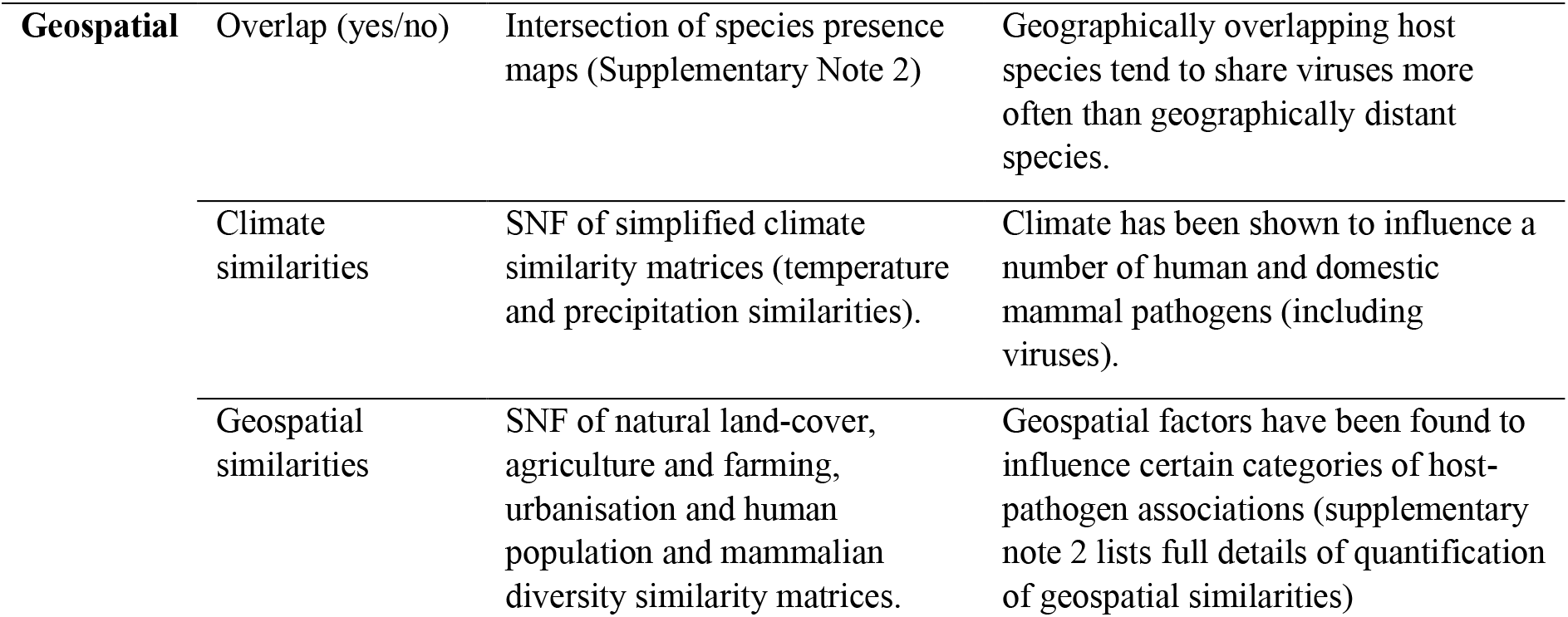
Mammalian phylogenetic, ecological, and geospatial similarities. Pairwise similarities were calculated for our mammalian species (n=876). Full details of these similarities, their sources and full justification are listed in Supplementary Note 2.

| Category | Similarities | Calculation | Reason for inclusion |
| --- | --- | --- | --- |
| **Phylogeny** | Phylogenetic similarity | 1-normalised phylogenetic distance. | Linked to sharing of viruses between mammals. |
| **Ecology** | Life-history and reproductive traits similarities. | Gower's distance matrices<sup>49</sup> | Life-history traits are a key feature in terms of metabolism and adaption to environment. Reproductive traits are potentially relevant in terms of within-host dynamics of viruses. |
|  | Habitat utilisation |  | Similar habitat utilisation might correlate with contact with similar viruses. |
|  | Diet (proportional use of 10 categories) |  | Similar dietary habit might associate with similar viral assemblage. |

| **Geospatial** | Overlap (yes/no) | Intersection of species presence maps (Supplementary Note 2) | Geographically overlapping host species tend to share viruses more often than geographically distant species. |
| --- | --- | --- | --- |
|  | Climate similarities | SNF of simplified climate similarity matrices (temperature and precipitation similarities). | Climate has been shown to influence a number of human and domestic mammal pathogens (including viruses). |
|  | Geospatial similarities | SNF of natural land-cover, agriculture and farming, urbanisation and human population and mammalian diversity similarity matrices. | Geospatial factors have been found to influence certain categories of host-pathogen associations (supplementary note 2 lists full details of quantification of geospatial similarities) |

Furthermore, we capture the changes in structure of the bipartite network linking coronaviruses with their mammalian hosts (figure 3). On one hand the nestedness of network increased (ranging from: 6.39-fold at 0.95 to 10.17-fold at 0.50 cut-off, table 1). On the other hand, the non-independence (C-score) of coronaviruses and mammalian hosts decreased with the addition of new links. Larger values of C-score suggest viral and host communities with little or no overlap in host or virus preferences (e.g. tendencies of coronaviruses to be shared amongst certain host communities, defined by phylogeny or geographical distribution), as visualised in figure 3.

**Figure 3.**
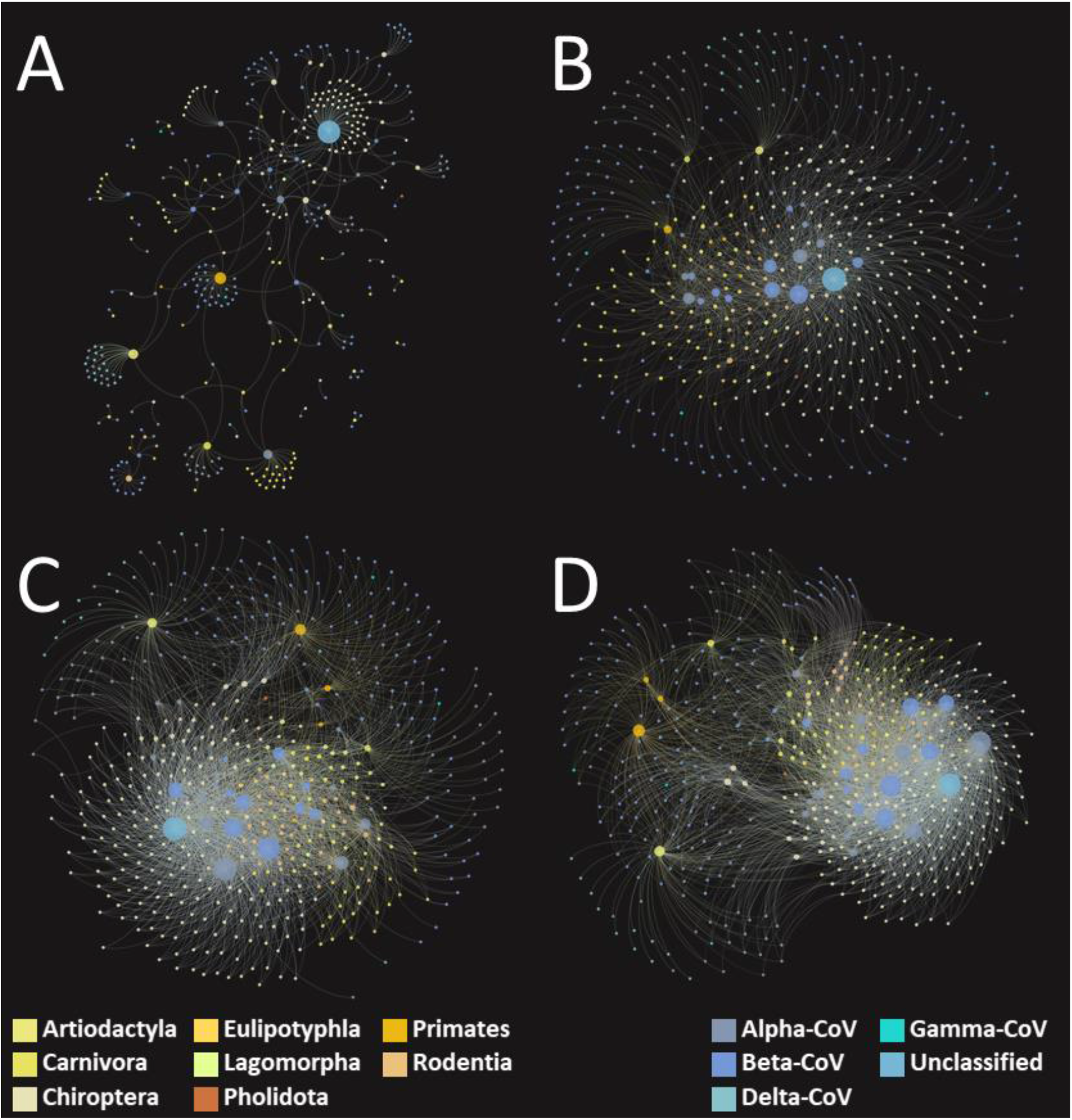
Bipartite networks linking coronaviruses with mammalian hosts. **Panel A: Original bipartite network** based on known/observed virus-host associations extracted from meta-data accompanying genomic sequences and supplemented with publications data from the Enhanced Infectious Diseases database (EID2). Panels B, C and D show predicted bipartite networks using our predicted virus-host associations at different cut-offs: 0.95, 0.75 and 0.5, respectively, for mean probability of associations.

### Validation

we validated our analytical pipeline externally against 5 held-out test-sets (as described in method section below). On average, our gbm ensemble achieved AUC = 0.95 (±0.028 SD), TSS=0.90 (±0.055), and F-Score = 0.10 (±0.068) (supplementary figure S6).

## DISCUSSION

In this study we deployed a meta-ensemble of similarity learners from three complementary perspectives (viral, mammalian and network), to predict the occurrence of associations between 411 known coronaviruses and 876 mammal species. We predict 4,438 new associations (11.54-fold increase), leading to the prediction that there are many more mammalian species than are currently known in which more than one coronavirus can occur. These hosts of multiple coronaviruses are potential sources of new coronavirus strains by homologous recombination. Here, we discuss the large number of candidate hosts in which homologous recombination of coronaviruses could result in the generation of novel pathogenic strains, as well as the substantial underestimation of the range of viruses which could recombine based on observed data. Our results are also discussed in terms of which host species are high priority targets for surveillance, both short and long-term.

Give that coronaviruses frequently undergo homologous recombination when they co-infect a host, and that SARS-CoV-2 is highly infectious to humans, the most immediate threat to public health is recombination of other coronaviruses with SARS-CoV-2. Such recombination could readily produce further novel viruses with both the infectivity of SARS-CoV-2 and additional pathogenicity or viral tropism from elsewhere in the Coronaviridae. Hence, in figure 1 (and supplementary results table SR1), we identify mammals predicted to be hosts of SARS-CoV-2 as well as multiple other coronaviruses. Taking only observed data, there are four non-human mammalian hosts known to associate with SARS-CoV-2 and at least one other coronavirus, and a total of 504 different unique interactions between SARS-CoV-2 and other coronaviruses (counting all combinations of virus and host individually). Any of these SARS-CoV-2 hosts which are also hosts of other coronaviruses are potential recombination hosts in which novel coronaviruses related to SARS-CoV-2 could be generated in future. However, when we add in our model’s predicted interactions this becomes 126 SARS-CoV-2 hosts and 2,544 and 1,302 total unique interactions, at 0.5 and 0.9 probability; indicating that observed data are missing 31.5-fold of the total number of potential recombination hosts and 5.05-fold increase (at 0.5 probability cut-off, 2.58-fold at 0.9) of the potential unique associations. These large-fold increases in the number of predicted hosts and associations demonstrate that the potential for homologous recombination between SARS-CoV-2 and other coronaviruses, which could lead to new pathogenic strains, is highly underestimated, both in terms of the range of hosts as well as the number of interactions within known hosts.

Our model has successfully highlighted known important recombination hosts of coronaviruses, adding confidence to our methodology. The Asian palm civet (*Paradoxurus hermaphroditus*), a viverrid native to south and southeast Asia, was identified by our model as a potential host of 32 and 20 different coronaviruses at 0.5 and 0.9 probability (vs 6 observed). Genetic evolution analysis has shown that SARS-CoV-2 is closely related to coronaviruses derived from *P. hermaphroditus*^26^, and its role as a reservoir for SARS-CoV^27^, strongly supporting our findings that it is an important host in viral recombination. This, together with the close association of *P. hermaphroditus*^26^ with humans, for example via bushmeat and the pet trade^28^ and in ‘battery cages’ for the production of Kopi luwak coffee^29^, highlights both the ability and opportunity of this species to act as a recombination host, with significantly more coronaviruses than have been observed. Furthermore, our model highlights both the greater horseshoe bat (*Rhinolophus ferrumequinum*), which is a known recombination host of SARS-CoV^30,31^, as well as the intermediate horseshoe bat (*Rhinolophus affinis*), which is believed to be recombination host of SARS-CoV-2^12,32^. Our model predicts *R. ferrumequinum* to be a host to 67 and 37 different coronaviruses at 0.5 and 0.9 probability respectively (vs 13 observed); and for *R. affinis* to host 44 and 22 at 0.5 and 0.9 probability respectively (vs 9 observed). Our model also successfully highlights the pangolin (*Manis javanica*), a suspected intermediate host for SARS-CoV-2^13^ as a potential host of 15 and 8 different coronaviruses at 0.5 and 0.9 probability (vs 1 observed).

The successful highlighting of the speculated main hosts for SARS-CoV and SARS-CoV-2 homologous recombination adds substantial confidence that our model is identifying the most important potential recombination hosts. Furthermore, our results show that the number of viruses which could potentially recombine even within these known hosts has been highly underestimated, indicating that there still remains significant potential for further novel coronavirus generation in future from current known recombination hosts.

In addition to highlighting known important recombination hosts, our model also identifies a diverse range of species not yet associated with SARS-CoV-2 recombination, but which are both predicted to host SARS-CoV-2 as well as large numbers of other coronaviruses. These hosts represent new targets for surveillance of novel human pathogenic coronaviruses. Amongst the highest priority is the lesser Asiatic yellow bat (*Scotophilus kuhlii*), a known coronavirus host^33^, common in east Asia but not well studied, and which features prominently with a large number of predicted interactions (47 at 0.5, 20 at 0.9). Figure 1 also implicates the common hedgehog (*Erinaceus europaeus*), the European rabbit (*Oryctolagus cuniculus*) and the domestic cat (*Felis catus*) as predicted hosts for SARS-CoV-2 (confirmed for the cat^34^) and large numbers of other coronaviruses (20, 23, 65 at 0.5; and 14, 14, 36 at 0.9, for the hedgehog, rabbit and cat, respectively). The hedgehog and rabbit have previously been confirmed as hosts for other betacoronaviruses^35,36^ which have no appreciable significance to human health. Our prediction of these species’ potential interaction with SARS-CoV-2 and considerable numbers of other coronaviruses, as well as the latter three species’ close association to humans, identify them as high priority underestimated risks. In addition to these human-associated species, both the chimpanzee (*Pan troglodytes*) and African green monkey (*Chlorocebus aethiops*) have large numbers of predicted associations (51, 46 at 0.5; and 16, 11 at 0.9, respectively), and given their relatedness to humans and their importance in the emergence of viruses such as DENV^37^ and HIV^38^, also serve as other high priority species for surveillance.

The most prominent result in figure 1 is the common pig (*Sus scrofa*), having the most predicted associations, in addition to SARS-CoV-2, of all included mammals (121 and 73 coronaviruses predicted at 0.5 and 0.9 cut-offs). The pig is a major known mammalian coronavirus host, harbouring both a large number (26) of observed coronaviruses, as well as a wide diversity (figure 2, supplementary results table SR4). Given the large number of predicted viral associations presented here, the pig’s close association to humans, its known reservoir status for many other zoonotic viruses^39^, and it being involved in genetic recombination of some of these viruses^40^, the pig is predicted to be one of the foremost candidates for an important recombination host.

In addition to the more immediate threat of homologous recombination directly with SARS-CoV-2, we also present our predicted associations between all mammals and all coronaviruses (figures 2 and 3). These associations represent the longer-term potential for background viral evolution via homologous recombination in all species. These data also show that there is a 11.54-fold underestimation in the number of associations, with 421 observed associations and 4438 predicted at 0.5 cut-off (5.72-fold increase with 1989 associations predicted at 0.9 cut-off). This is visually represented in figure 3, which shows the bipartite network of virus and host for observed associations (A), and predicted associations (B-D); with a marked increase in connectivity between our mammalian hosts and coronaviruses, even at the most stringent 0.95 confidence cut-off. This indicates that the potential for homologous recombination between coronaviruses is substantially underestimated using just observed data.

Furthermore, our model shows that the associations between more diverse coronaviruses is also underestimated, for example, the number of host species with four or more different subgenera of coronaviruses increases by 41.57-fold from seven observed to 291 predicted (Table S4, shows the degree of diversity of coronaviruses in mammalian host species highlighted in figure 2). The high degree of potential co-infections including different sub-genera and genera seen in our results emphasises the level of new genetic diversity possible via homologous recombination in these host species (table S4). A similar array of host species is highlighted for total associations as was seen for SARS-CoV-2 potential recombination hosts, including the common pig, the lesser Asiatic yellow bat, and both the greater and intermediate horseshoe bats, whilst notable additions include the dromedary camel (*Camelus dromedaries*). The camel is a known host of multiple coronaviruses and the primary route of transmission of MERS-CoV to humans^41,42^. Our results suggest that monitoring for background viral evolution via homologous recombination would focus on a similar array of hosts, with a few additions, as monitoring for SARS-CoV-2 recombination. Again, our results strongly suggest that the potential array of viruses which could recombine in hosts is substantially underestimated, reinforcing the message that continued monitoring is essential.

Methodologically, the novelty of our approach lies in integrating three points of view: that of the coronaviruses, that of their potential mammalian hosts, and that of the network summarising our knowledge to date of sharing of coronaviruses in their hosts. By constructing a comprehensive set of similarity learners in each point of view and combining these learners non-linearly (via GBM meta-ensemble) a great strength of our analytical pipeline is that it is able to predict potential recombination hosts of coronaviruses without any prerequisite knowledge or assumptions into which parts of the coronavirus genomes are important, or integration of receptor information, or focusing on certain groups of hosts (e.g. bats or primates). This ‘no-preconceptions’ approach enables us to analyse without being restricted by our current incomplete knowledge of the specific biological and molecular mechanisms which govern host-virus permissibility Additionally, the incorporation of similarity-based learners in our three-perspective approach enabled us to capture new hosts (i.e. with no known association with coronaviruses), thus avoiding a main limitation of approaches which rely only on networks and their topology.

We acknowledge certain limitations in our methodology, primarily pertaining to current incomplete datasets in the rapidly developing but still understudied field. 1) The inclusion only of coronaviruses for which complete genomes could be found limited the number of coronaviruses (species or strain) for which we could compute meaningful similarities, and therefore predict potential hosts. The same applies for our mammalian species – we only included mammalian hosts for which phylogenetic, ecological, and geospatial data were available. As more data on sequenced coronaviruses or mammals become available in future, our model can be re-run to further improve predictions, and to validate predictions from earlier iterations. 2) Research effort, centering mainly on coronaviruses found in humans and their domesticated animals, can lead to overestimation of the potential of coronaviruses to recombine in frequently studied mammals, such as lab rodents which were excluded from the results reported here (similar to previous work^18^), and significantly, domesticated pigs and cats which we have found to be important recombination host species of coronaviruses. This latter limitation is partially mitigated. Firstly, in our results, other ‘overstudied’ mammals such as cows and sheep, were not highlight by our model, which is consistent with them being considered less important hosts of coronaviruses, and certain under-studied bats were highlighted as major potential hosts; together, these indicate that research effort is not a substantial driver of our results. Secondly, methodologically, the effect of research effort has been mitigated by capturing similarities from our three points of view (virus, host and network) and multiple characteristics therein.

In this study we demonstrate that the potential for homologous recombination in mammalian hosts of coronaviruses is highly underestimated. The ability of the large numbers of hosts presented here to be hosts of multiple coronaviruses, including SARS-CoV-2, demonstrated the capacity for homologous recombination and hence production of further novel coronaviruses. Our methods deployed a meta-ensemble of similarity learners from three complementary perspectives (viral, mammalian and network), to predict each potential coronavirus-mammal association.

The current consensus is that SARS-CoV-2 was generated by homologous recombination; originally derived from coronaviruses in bats^12^ and then shifted to humans via an intermediate reservoir host, likely a species of pangolin^13^. Importantly, the lineage of SARS-CoV-2 was deduced only after the outbreak in humans. With the greater understanding of the extent of mammalian host reservoirs and the potential recombination hosts we identify here, a targeted surveillance program is now possible which would allow for this generation to be observed as it is happening and before a major outbreak. Such information could help inform mitigation strategies and provide a vital early warning system for future novel coronaviruses.

## METHODS

### Viruses and mammalian data

#### Viral genomic data

Complete sequences of coronaviruses were downloaded from Genbank^43^. Sequences labelled with the terms: “vaccine”, “construct”, “vector”, “recombinant” were removed from the analyses. In addition, we removed these associated with experimental infections were possible. This resulted in total of 3,264 sequences for 411coronavirus species or strain (i.e. viruses below species level on NCBI taxonomy tree). Of these 88 were sequences of coronavirus species, and 307 of strains (in 25 coronavirus species, with total number of species included=92).

### Selection of potential mammalian hosts of coronaviruses

We processed meta-data accompanying all sequences (including partial sequences but excluding vaccination and experimental infections) of coronaviruses uploaded to GenBank to extract information on hosts (to species level) of these coronaviruses. We supplemented these data species-level hosts of coronaviruses extracted from scientific publications via the Enhanced Infectious Diseases Database (EID2)44. This resulted in identification of 313 known terrestrial mammalian hosts of coronaviruses (regardless of whether a complete genome was available or not, N=185 mammalian species for which an association with a coronaviruses with complete genome was identified). We expanded this set of potential hosts by including terrestrial mammalian species in genera containing at least one known host of coronavirus, and which are known to host one or more other virus species (excluding coronaviruses, information of whether the host is associated with a virus were obtained from EID2). This results in total of 876 mammalian species which were selected.

### Quantification of viral and mammalian similarities

We computed three types of similarities between each two viral genomes as summarised below.

#### Biases and codon usage

We calculated proportion of each nucleotide of the total coding sequence length. We computed dinucleotide and codon biases^45^ and codon pair bias, measured as the codon pair score (CPS)^45,46^ in each of the above sequences. This enabled us to produce for each genome sequence (N=3,271) the following feature vectors: nucleotide bias; dinucleotide bias; codon biased; and codon-pair bias.

#### Secondary structure

Following alignment of sequences (using *AlignSeqs* function in R package *Decipher*^47^), we predicted the secondary structure for each sequence using *PredictHEC* function in the R package *Decipher*^47^. We obtained both states (final prediction), and probability of secondary structures for each sequence. We then computed for each 1% of the genome length both the coverage (number of times a structure was predicted) and mean probability of the structure (in the percent of the genome considered). This enabled us to generate six vectors (length = 100) for each genome representing: mean probability and coverage for each of three possible structures – Helix (H), Beta-Sheet (E), or Coil (C).

#### Genome dissimilarity (distance)

we calculated pairwise dissimilarity (in effect a hamming distance) between each two sequences in our set using the function *DistanceMatrix* in the R package *Decipher*^47^. We set this function to penalise gap-to-gap and gap-to-letter mismatches.

#### Similarity quantification

We transformed the feature (traits) vectors described above into similarities matrices between coronaviruses (species or strains). This was achieved by computing cosine similarity between these vectors in each category (e.g. codon pair usage, H coverage, E probability). Formally, for each genomic feature (N=10) presented by vector as described above, this similarity was calculated as follows:

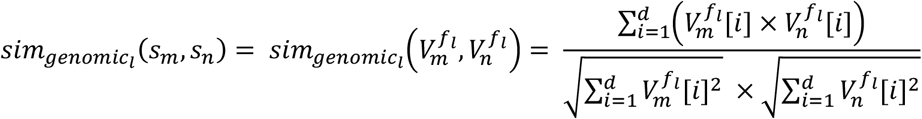

Where *s*_*m*_ and *s*_*n*_ are two genomic sequences presented by two feature vectors 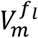 and 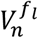 from the genomic feature space *f*_*l*_ (e.g. codon pair bias) of the dimension d (e.g. d= 100 for H-coverage).

We then calculated similarity between each pair of virus strains or species (in each category) as the mean of similarities between genomic sequences of the two virus strains or species (e.g. mean nucleotide bias similarity between all sequences of SARS-CoV-2 and all sequences of MERS-CoV presented the final nucleotide bias similarity between SARS-CoV-2 and MERS-CoV). This enabled us to generate 11 genomic features similarity matrices (the above 10 features represented by vectors + genomic similarity matrix) between our input coronaviruses.

#### Similarity network fusion (SNF)

We applied similarity network fusion (SNF)^48^ to integrate the following similarities in order to reduce our viral genomic feature space: 1) Nucleotide, dinucleotide, codon, and codon pair usage biases were combined into one similarity matrix - *genome bias similarity*. And 2) Helix (H), Beta-Sheet (E), or Coil (C) mean probability and coverage similarities (six in total) were combined into one similarity matrix - *secondary structure similarity*.

SNF applies an iterative nonlinear method that updates every similarity matrix according to the other matrices via nearest neighbour approach and is scalable and is robust to noise and data heterogeneity. The integrated matrix captures both shared and complementary information from multiple similarities.

#### Mammalian similarities

We calculated comprehensive set of mammalian similarities. Table 1 summarises these similarities and provides justification for inclusion. Supplementary note 2 provides full details.

### Quantification of network similarities

#### Network construction

We processed meta-data accompanying all sequences (including partial genome but excluding vaccination and experimental infections) of coronaviruses uploaded to GenBank to extract information on hosts (to species level) of these coronaviruses. We supplemented these data with virus-host associations extracted from publications via the Enhanced Infectious Diseases Database (EID2)^44^. This resulted in 1,669 associations between 1108 coronaviruses and 545 hosts (including non-mammalian hosts). We transformed this set of associations into a bipartite network linking species and strains of coronaviruses with their hosts.

#### Quantification of topological features

The above constructed network summarises our knowledge to date of associations between coronaviruses and their hosts, and its topology expresses patterns of sharing these viruses between various hosts and host groups. Our analytical pipeline captures this topology, and relations between nodes in our network, by means of node (vertex) embeddings. This approach encodes each node (here either a coronavirus or a host) with its own vector representation in a continuous vector space, which in turns enable us to calculate similarities between two nodes based on this representation.

We adopted DeepWalk^23^ to compute vectorised representations for our coronaviruses and hosts from the network connecting them. DeepWalk uses truncated random walks to get latent topological information of the network and obtains the vector representation of its nodes (in our case coronaviruses, and their hosts) by maximizing the probability of reaching a next node (i.e. probability of a virus-host association) given the previous nodes in these walks (Supplementary Note 3 provides formal and visual summaries of DeepWalk).

#### Similarity calculations

Following the application of DeepWalk to compute the latent topological representation of our nodes, we calculated the similarity between two nodes in our network - n (vectorised as N) and m (vectorised as M), by using cosine similarity as follows ^24,25^:

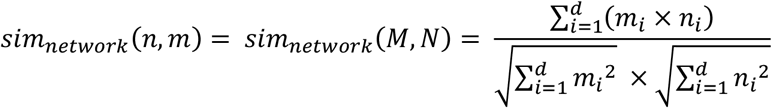

where d is the dimension of the vectorised representation of our nodes: M and N, and *m*_*i*_ and *n*_*i*_ are the components of vectors M and N, respectively.

### Similarity learning meta-ensemble – a multi-perspective approach

Our analytical pipeline stacks 12 similarity learners into meta-ensembles and selects the best performing ensemble following internal and external validation processes. The constituent learners based on different data and points of views and can be categorised as follows:

***Coronaviruses – the viral point of view***: we assembled three models derived from a) genome similarity, b) genome biases, and c) genome secondary structure. Each of these learners gave each coronavirus-mammalian association (*v*_*i*_ → *m*_*j*_) a score, termed confidence, based on how similar the coronavirus *v*_*i*_ is to known coronaviruses of mammalian species *m*_*j*_, compared to how similar *v*_*i*_ is to all included coronaviruses^24,25^. In other words, if *v*_*i*_ is more similar (e.g. based on genome secondary structure) to coronaviruses observed in host *m*_*j*_ than it is similar to those coronaviruses not observed in *m*_*j*_, then the association *v*_*i*_ → *m*_*j*_ is given a higher confidence score. Conversely, if *v*_*i*_ is somewhat similar to coronaviruses observed in *m*_*j*_, and also somewhat similar to viruses not known to infect this particular mammal, then the association *v*_*i*_ → *m*_*j*_ is given a medium confidence score. The association *v*_*i*_ → *m*_*j*_ is given a lower confidence score if *v*_*i*_ is more similar to coronaviruses not known to infect *m*_*j*_ than it is similar to coronaviruses observed in this host.

Formally, given an adjacency matrix A of dimensions |*V*| × |*M*| where |*V*| is number of coronaviruses included in this study (for which a complete genome could be found), and |*M*| is number of included mammals, such that for each *v*_*i*_ ∈ *V* and *m*_*j*_ ∈ *M*, *a*_*ij*_ = 1 if an association is known to exist between the virus and the mammal, and 0 otherwise. Then for a similarity matrix *sim*_*viral*_ corresponding to each of the similarity matrices calculated above, a learner from the viral point of view is defined as follows^24,25^:

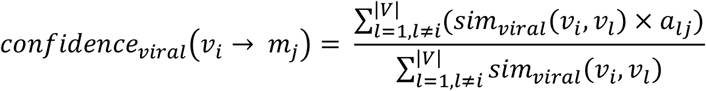

***Mammals – the host point of view***: we constructed 7 learners from the similarities summarised in table 1. Each of these learners calculated for every coronavirus-mammalian association (*v*_*i*_ → *m*_*j*_) a confidence score based on how similar the mammalian species *m*_*j*_ is to known hosts of the coronavirus *v*_*i*_, compared to how similar *m*_*j*_ is to mammals not associated with *v*_*i*_. For instance, if *m*_*j*_ is phylogenetically close to known hosts of *v*_*i*_, and also phylogenetically distant to mammalian species not known to be associated with this coronavirus then the phylogenetic similarly learner will assign *v*_*i*_ → *m*_*j*_ a higher confidence score. However, if *m*_*j*_ does not overlap geographically with known hosts of *v*_*i*_, then the geographical overlap learner will assign it a low (in effect 0) confidence score.

Formally, given the above defined adjacency matrix A, and a similarity matrix *sim*_*mammalian*_ corresponding to each of the similarity matrices summarised in table 1 and calculated in Supplementary Note 2, a learner from the mammalian point of view is defined as follows^24,25^:

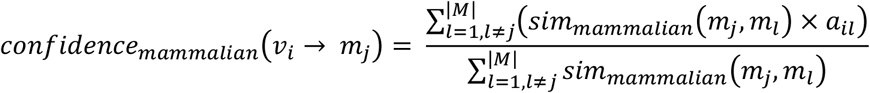

***Network – the network point of view***: we integrated two learners based on network similarities - one for mammals and one for coronaviruses. Formally, given the adjacency matrix A, our two learners from the network point of view as defined as follows ^24^:

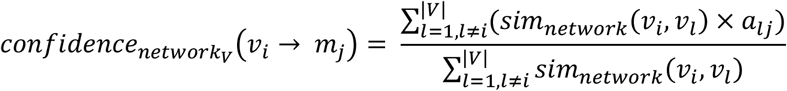

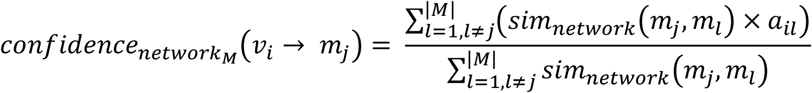

### Ensemble construction

We combined the above described leaners by stacking them into ensembles (meta-ensembles) using Stochastic Gradient Boosting (GBM). The purpose of this combination is to incorporate the three points of views descried above, as well as varied aspects of the coronaviruses and their mammalian potential hosts, into a generalisable, robust model^50^. We selected GBM as our stacking algorithm due to its ability to handle non-linearity and high-order interactions between constituent learners^51^. We performed the training and optimisation (tuning) of these ensembles using the *caret R Package*^52^.

### Ensemble sampling

our gbm ensembles comprised 100 replicate models. Each model was trained with balanced random samples using 10-fold cross validation. Final ensembles were generated by taking mean predictions (probability) of constituent models. Standard deviation (SD) from mean probability was also generated and its values subtracted/added to predictions.

### Validation and performance estimation

We validated the performance of our analytical pipeline externally against 5 held-out test-sets. each test-set was generated by splitting the set of observed associations between coronaviruses and their hosts into two random sets: a training-set comprising 85% of all known associations and a test-set comprising 15% of known associations. These test-sets were held-out throughout the processes of generating similarity matrices; similarity learning, and assembling our learners, and were only used for the purposes of estimating performance metrics of our analytical pipeline. This resulted in 5 runs in which our ensemble learnt using only 85% of observed associations between our coronaviruses and their mammalian hosts. For each run calculated three performance metrics based on the mean probability across each set of 100 replicate models of the gbm meta-ensembles: AUC, true skills statistics (TSS) and F-score.

AUC is a threshold-independent measure of model predictive performance that is commonly used as a validation metric for host-pathogen predictive models^21,45^. Use of AUC has been criticised for its insensitivity to absolute predicted probability and its inclusion of a priori untenable prediction^51,53^, we also calculated the True Skill Statistic (TSS = sensitivity + specificity −1)^54^. F-score captures the harmonic mean of the precision and recall and is often used with uneven class distribution. Our approach is relaxed with respect to false positives s(unobserved associations), hence the low F-score recorded overall.

### Changes in network structure

We quantified the diversity of the mammalian hosts of each CoV in our input by computing mean phylogenetic distance between these hosts. Similarly, we captured the diversity of CoVs associated with each mammalian species by calculating mean (hamming) distance between the genomes of these CoVs. We termed these two metrics: mammalian diversity per virus and viral diversity per mammal, respectively. We aggregated both metrics at the network level by means of simple average. This enabled us to quantify changes in these diversity metrics, at the level of network, with addition of predicted links at three probability cut-offs: >0.5, >0.75 and ≥0.95.

In addition, we captured changes in the structure of the bipartite network linking CoVs with their mammalian hosts, with the addition of predicted associations, by computing comprehensive set of structural properties (Supplementary Note 4) at the probability cut-offs mentioned above, and compare the results with our original network. Here we ignore properties that deterministically change with addition of links (e.g. degree centrality, connectance, table S2 lists all computed metrics and changes in their values). Instead we focus on non-trivial structural properties. Specifically, we capture changes in network stability, by measuring its nestedness^55–57^; and we quantify non-independence in interaction patterns by means of Checkerboard score^58^ (C-Score). Supplementary note 4 provides full definition of these concepts as well as other metrics we computed for our networks.

## Supporting information

Supplementary results table - SR1

Supplementary results table - SR2

Supplementary results table - SR3

Supplementary results table - SR4

## ACKNOWLEDGMENTS

MW acknowledges support from BBSRC and MRC for the National Productivity Investment Fund (NPIF) fellowship (MR/R024898/1). Establishment of the EID2 database was funded by a UK Research Council Grant (NE/G002827/1) to MB, as part of an ERANET Environmental Health award to MB; subsequently, it has been further developed and maintained by BBSRC Tools and Resources Development Fund awards (BB/K003798/1; BB/N02320X/1) to MB, and the National Institute for Health Research Health Protection Research Unit (NIHR HPRU) in Emerging and Zoonotic Infections at the University of Liverpool in partnership with Public Health England and Liverpool School of Tropical Medicine.

## AUTHOR CONTRIBUTIONS

Conceived and designed the study: MW and MSCB

Compiled the data: MW

Analysed and interpreted the data: MW and MSCB

Established the EID2 database: MW and MB

Wrote the manuscript: MW, MB and MSCB.

## MATERIALS & CORRESPONDENCE

Maya Wardeh and Marcus SC Blagrove

## DATA AVAILABILITY STATEMENT

All data used in our analyses will be made available via figshare. During review process our data can be made available upon request.

## CODE AVAILABILITY STATEMENT

All codes used in our analyses will be made available via figshare. During review process our code can be made available upon request.

