## Supplementary notes for "Predicting mammalian hosts in which novel coronaviruses can be generated"

### Electronic Supplementary Materials

**Maya Wardeh*, Matthew Baylis, Marcus SC Blagrove***

#### Supplementary Note 1 – From viral genomic traits to viral similarity

Complete sequences of coronaviruses were downloaded from Genbank [1,2]. Sequences labelled with the terms: "vaccine", "construct", "vector", "recombinant" were removed from the analyses. In addition, we removed these associated with experimental infections were possible. This resulted in total of **3,168** sequences for **395** coronavirus species (N=88) or strain (i.e. viruses below species level on NCBI taxonomy tree, N=307, in 25 coronavirus species, with total number of species included=92).

##### Biases and codon usage

We calculated proportion of each nucleotide [A, C, G, T] of the total coding sequence length. We computed dinucleotide and codon biases [3] and codon pair bias, measured as the codon pair score (CPS) [3,4] in each of the above sequences. This enabled us to produce for each genome sequence (N=3,168) the following feature vectors:

1. Nucleotide bias: 4 features.
2. Dinucleotide bias: 16 features.
3. Codon bias: 64 features.
4. Codon pair bias: 3,721features.

##### Secondary structure

Following alignment of sequences, we predicted the secondary structure for each sequence using *PredictHEC* function in the R package *Decipher*[5]. We obtained both states (final prediction), and probability of secondary structures for each sequence. We then computed for each 1% of the genome length both the coverage (number of times a structure was predicted) and mean probability of the structure (in the percent of the genome considered). This enabled us to generate six vectors (length = 100) for each genome representing: mean probability and coverage for each of three possible structures – Helix (H), Beta-Sheet (E), or Coil (C).

##### Genome dissimilarity (distance)

Following alignment of sequences, we calculated pairwise dissimilarity (in effect a hamming distance) between each two sequences in our set using the function *DistanceMatrix* in the R package *Decipher* [5]. We set this function to penalise gap-to-gap and gap-to-letter mismatches.

##### Similarity quantification

We transformed the feature (traits) vectors calculated above into similarities matrices between coronaviruses (species or strains). This was done by computing cosine similarity between these vectors in each category (e.g. codon pair usage, H coverage, E probability).
by computing cosine similarity between vectors of features in each category (e.g. codon pair usage, H coverage, E probability). Formally, for each genomic feature (N=10) presented by vector as described above, this similarity was calculated as follows:

$$sim_{genomic_l}(s_m, s_n) = sim_{genomic_l}\left(V_m^{f_l}, V_n^{f_l}\right) = \frac{\sum_{i=1}^{d}\left(V_m^{f_l}[i] \times V_n^{f_l}[i]\right)}{\sqrt{\sum_{i=1}^{d} V_m^{f_l}[i]^2} \times \sqrt{\sum_{i=1}^{d} V_n^{f_l}[i]^2}}$$

Where $s_m$ and $s_n$ are two genomic sequences presented by two feature vectors $V_m^{f_l}$ and $V_m^{f_l}$ from the genomic feature space $f_l$ (e.g. codon pair bias) of the dimension d (e.g. d= 3,721 for codon pair bias).

We then calculated similarity between each pair of virus strains or species (in each category) as the mean of similarities between genomic sequences of the two virus strains or species (e.g. mean nucleotide bias similarity between all sequences of SARS-CoV-2 and all sequences of MERS- CoV presented the final nucleotide bias similarity between SARS-CoV-2 and MERS-CoV). This enabled us to generate 11 genomic features similarity matrices (the above 10 features represented by vectors + genomic dissimilarity matrix) between our input coronaviruses. Figure S1 illustrate the process.

#### Similarity network fusion (SNF):

We applied similarity network fusion (SNF) [6], via the R package *SNF*, to integrate the following similarities (computed above) in order to reduce our viral genomic feature space:

1. Nucleotide, dinucleotide, codon, and codon pair usage biases were combined into one similarity matrix - *genome bias similarity*.
2. Helix (H), Beta-Sheet (E), or Coil (C) mean probability and coverage similarities (six in total) were combined into one similarity matrix - *secondary structure similarity*.

SNF applies an iterative nonlinear method that updates every similarity matrix according to the other matrices via nearest neighbour approach (KNN) and is scalable and is robust to noise and data heterogeneity. The integrated similarity matrix captures both shared and complementary information from multiple similarities.

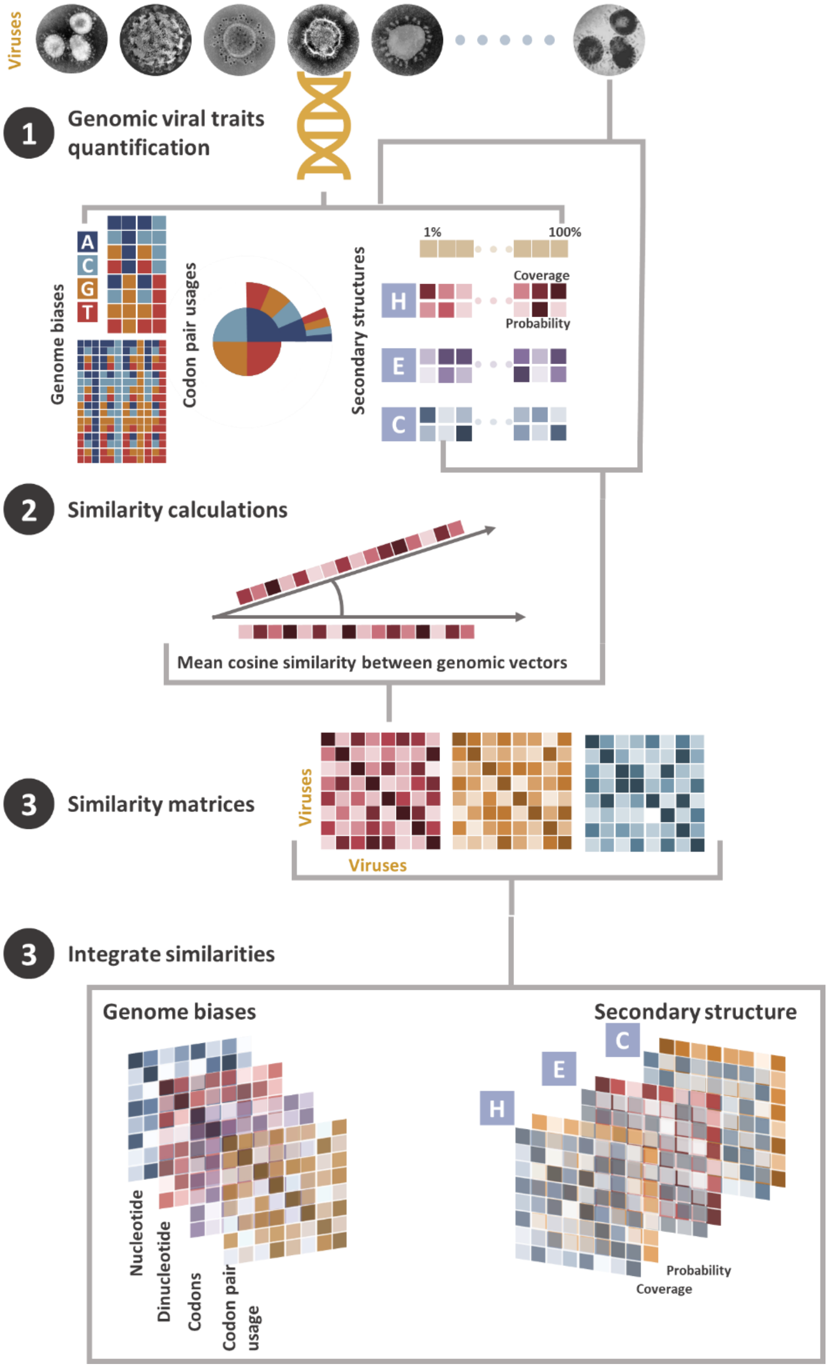


**Figure S1 – Computing genomic-features similarity matrices of coronaviruses.** First genomic traits were extracted from complete genome sequences of coronaviruses– step 1. Then the vector representations of each trait were used to compute cosine similarity between two genomes for the trait (e.g. dinucleotide biases) and the mean was taken when computing similarity between two coronaviruses represented by one or more genome each – step 2. The resulting similarity matrices were integrated using SNF[6] to generate two integrated similarity matrices: *genome biases similarity* and *secondary structure similarity*.

#### Supplementary Note 2 – From mammalian phylogenetic, ecological and geo-spatial traits to mammalian similarity

Selection of potential mammalian hosts of coronaviruses:

We processed meta-data accompanying all sequences (including partial sequences but excluding vaccination and experimental infections) of coronaviruses uploaded to GenBank to extract information on hosts (to species level) of these coronaviruses. We supplemented these data species-level hosts of coronaviruses extracted from scientific publications via the Enhanced Infectious Diseases Database (EID2)[7]. This resulted in identification of 313 known terrestrial mammalian hosts of coronaviruses.

Following this, we expanded our set of potential hosts by including terrestrial mammalian species in genera containing a at least one known host of coronavirus, and which are known to host one or more other virus species (excluding coronaviruses, information of whether the host is associated with a virus was also extracted from EID2). This results in total of 876 mammalian species which were selected.

Phylogeny:

Mammalian phylogenetic distance has been linked to sharing of viruses[8–10]. We calculated pair wise phylogenetic similarity between each mammal-mammal pair based on phylogenetic distances extracted from a recent mammalian supertree[11].

Ecological traits:

We compiled data on morphological and life-history traits, diet and habitat for our mammal species from online databases and literature[12–17]. We selected the following traits for their known correlation with host-pathogen associations, and their wide availability: Body mass (g), maximum age (months), proxied key features of metabolism and adaption to environment; activity cycle, and migration[13] presented key aspects of mammalian behaviour. We utilised the following reproductive traits: age at sexual maturity (days), gestation period length (days), litters per year, litter size and weaning age (days). Reproductive traits could be viewed as proxies to within-host virus-dynamics and therefore may influence the viruses harboured by the host.

We incorporated the above traits to calculate traits-based pairwise distance between each two mammalian species. We based these distance calculations a generalised form of Gower's distance matrices [18,19]. We then transformed these distances into similarities (similarity between two mammals = 1- normalised distance).

Mammals utilising similar habitats might encounter similar coronaviruses and this in turn would increase the chances of being infected with these coronaviruses. We therefore incorporated habitat utilisation[16] as multiple binary indicators of whether a species uses one or more of 14 natural and artificial habitats. We transformed these habitat utilisations into similarity matrix following same procedure as above.

In addition to habitat, similar diet preference, expressed in terms of proportional use of 10 diet categories[15], could potentially associate with similar viral assemblage. We transformed these diet categories into a similarity matrix as per above.

The above steps we resulted in the following pair-wise ecological similarities between each pair of mammals in our study:

1. *Life-history and reproductive traits similarity*
2. *Habitat utilisation similarity*
3. *Diet similarity*

Geospatial traits:

The geographical distribution of mammalian species influences the coronaviruses with which they might come into contact. Geographical spread correlates with other factors such as climate, natural environment, and agricultural practices (including potential contact with livestock). Climate has been shown to influence a number of human and domestic mammal pathogens (including viruses)[20–22]. Other geographical factors such as biodiversity (species richness), land cover type, agriculture and farming practices, urbanisation and human population have been found to influence certain categories of host-pathogen associations[23,24].

**Species-presence maps**: We obtained species-presence maps for majority of our mammalian species from IUCN[16]. We extrapolated livestock (including horses) species-presence maps from most recent global distribution maps [25]. Finally, we inferred presence-maps for three domesticated species - dogs (*Canis lupus familiaris*), cats (*Felis catus)* and guinea pigs (*Cavia porcellus*) from Gridded population of the world maps [26], by assuming they co-exist with humans where there is sufficient human populations (n>100). We used the same gridded population maps to extrapolate human species-presence map (n>0). All our geographical maps manipulation was done in QGIS.

**Presence overlap**: we intersected the above curated maps using the R Package *raster* to compute whether the presence of any two mammalian species in our input overlapped (binary, 1=yes, 0=no), and to calculate the area of this overlap (in $km^2$).

**Vectorised geospatial features:** We supplemented species-presence maps with grids expressing climate[27], mammalian diversity [28], human population [26], land cover (including urbanisation) [29], agriculture[29,30], and distribution of livestock [25]. This allowed to generate the following geospatial feature vectors for each mammalian species (figure S2):

1. *Climate*: we expressed climate in two *vectorised features* as follows:
    a. Mean temperature: we computed mean of monthly temperatures recorded in each grid (table S1) in the species-presence area, averaged between years: 1900-2010[27]. WE transformed this gridded temperature in to an 11 points quantile vector representing the probabilities: 5%, 10%, 20%, 30%, 40%, 50%, 60%, 70%, 80%, 90%, and 95%.
    b. Mean precipitation: Sum of monthly rainfall (precipitation) recorded in each grid in the species-presence area, averaged between years: 1900-2010[27]. Gridded precipitation was transformed into quantile vector as above.
2. *Natural land-cover type* (not directly associated with humans): we computed vectorised features (see above) for each of the land-cover types in this category (table S1).
3. *Agricultural land-cover type* (including land-cover associated with humans e.g. managed vegetation) and *farming* practices (expressed in number of domesticated livestock and poultry in the species presence area) were also quantified into vectorised features as above.
4. *Urbanisation and human population*[26] vectorised features were computed form species-presence maps.
5. M*ammalian diversity*[28] in the species presence area was transformed into vectorised features.

We transformed the above vectors into pair-wise mammal-mammal similarity matrices by calculating cosine similarity between the vectorised feature (as per Supplementary Note 1).

Similarity network fusion (SNF):

We applied similarity network fusion (SNF)[6] to integrate the following similarity matrices calculated above in order to reduce our mammalian feature space:

1. *Climate* similarities: temperature and precipitation similarities were integrated using SNF.
2. *Geo-spatial traits*: the 7 similarities derived from natural land-cover type (table S1), the 11 11 similarities derived from agricultural land-cove type and farming (livestock and poultry); the 2

similarities based on urbanisation and human population, and the mammalian diversity similarity were integrated using SNF.

**Table S1 - List of geographical predictor layers integrated within our framework.**

| Category | layers(s)/geo-attributes | Source | Res | Reason |
|---|---|---|---|---|
| **(Natural) Land-cover** | Evergreen/deciduous needle-leaf trees (%) | EarthEnv [29] | 0°0′30″ | Type of land cover has been associated with distribution of various mammals [31]. It potentially increases chances of contact between mammalian reservoirs of different viruses. |
| | Evergreen broad-leaf trees (%) | | | |
| | Deciduous broad-leaf trees (%) | | | |
| | Mixed/other trees (%) | | | |
| | Shrubs (%) | | | |
| | Herbaceous vegetation (%) | | | |
| | Barren land (%) | | | |
| **Agriculture & farming** | Managed/Cultivated Vegetation (%) | EarthEnv [29] | 0°0′30″ | |
| | Regularly flooded vegetation (%) | | | |
| | Cropland (%) | HYDE [32] | 0°5′ | |
| | Pasture (%) | | | |
| | Cattle (head count) | [25] | 0.0833° | Livestock farming is linked to cross-species transmission of number of viruses[33]. |
| | Sheep (head count) | | | |
| | Buffalo (head count) | | | |
| | Pigs (head count) | | | |
| | Horses (head count) | | | |
| | Chicken (head count) | | | |
| | Duck (head count) | | | |
| **Human** | Human population | SEDAC [26] | 0°5′ | Urbanisation and human population density have been shown to be drivers of viral spill-over through wildlife-domestic-human interface[24,34,35]. |
| | Urban land (%) | EarthEnv [29] | 0°0′30″ | |
| **Climate** | Mean temperature | CRUTS3 [27] | 0°5′ | Climate could potentially influence the spread and emergence of viruses[20–22]. |
| | Mean precipitation | | | |
| **Mammalian diversity** | Number of different mammalian species in a grid cell. | SEDAC [28] | 0°5′ | Mammalian species present in mammal rich areas might be exposed to diverse viruses[36–39]. |

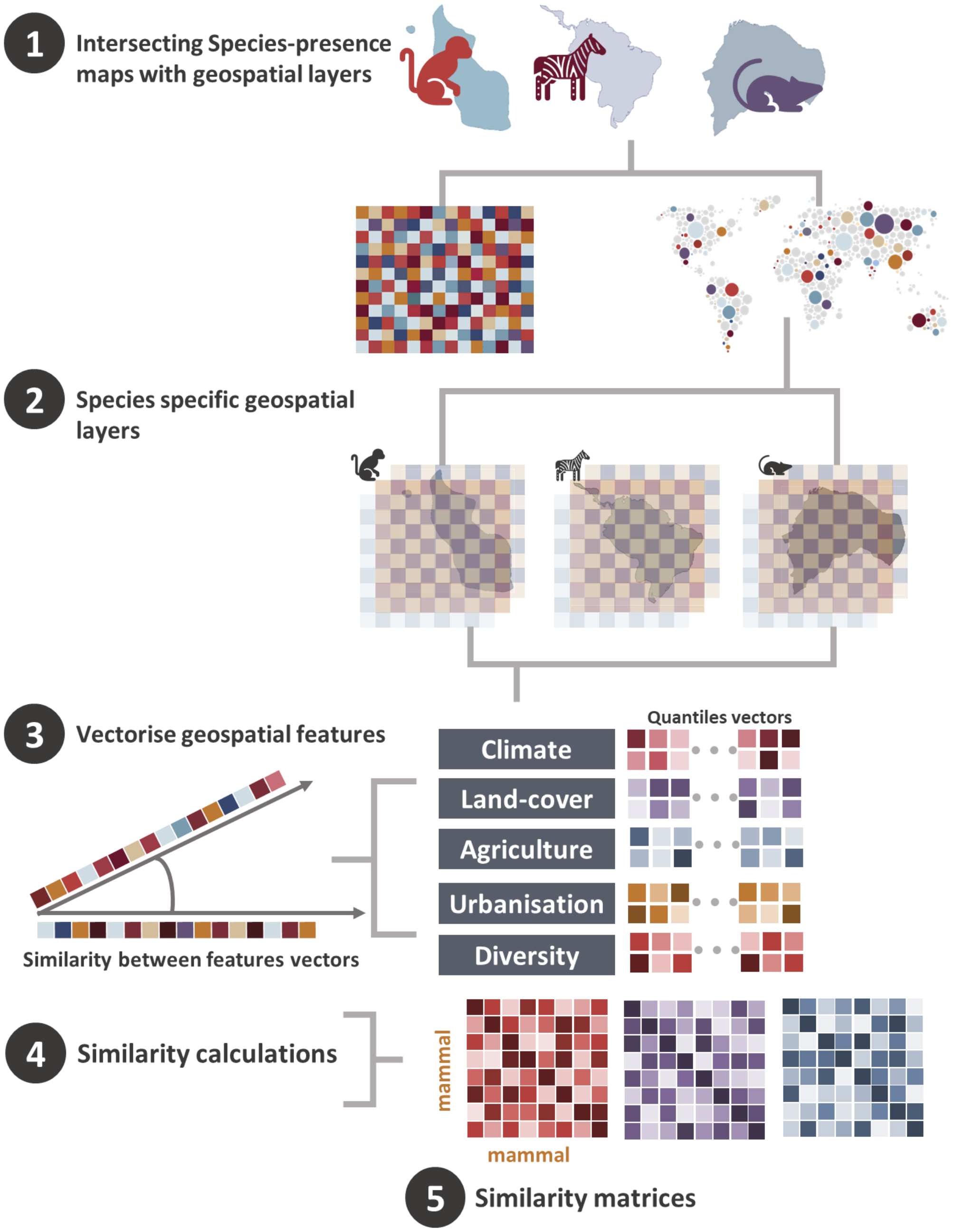


**Figure S2 – Vectorised geospatial features extraction**. Mammalian species-presence maps were first extracted from our sources[16,25,26]; these maps were then intersected with our geospatial layers (table S1) – step1. This enabled us to derive quantile vectors of our geospatial attributes (table S1) for majority of our mammalian species – step2, which we then transformed into vectors expressing quantile distribution of these attributes in the species presence area – step 3. Finally, we computed cosine similarity between these vectors to generate a pair-wise similarity matrix (between mammalian species) for each of our geospatial attributes – step 4.

#### Supplementary Note 3 – Network similarity

We adopted DeepWalk [40] to compute vectorised representations for our coronaviruses and hosts from the network connecting them. DeepWalk uses truncated random walks to get latent topological information of the network and obtains the vector representation of its nodes (in our case coronaviruses, and their hosts) by maximizing the probability of reaching a next node (i.e. probability of a virus-host association) given the previous nodes in these walks.

DeepWalk comprises three steps (figure S3):

1. *Sampling:* For each node $n_i$ (virus or host) in our network, DeepWalk conducts $\gamma$ random walks with length *t* starting from$n_i$*i.*
2. *Training skip-gram:* by treating walks as the equivalent of sentences, DeepWalk updates the node representation using the skip-gram algorithm [41] for each walk (figure S3). Here, skip-gram is used to maximise the cooccurrence probability among nodes which appear within a window *w* using an independent assumption as follows:

$$\Pr\left(\{n_{i-w}, \dots, n_{i+w}\} \backslash n_i | \phi(n_i)\right) = \prod_{j=i-w, j\neq i}^{i+w} \Pr\left(n_i | \phi(n_i)\right)$$

   where Φ denotes the latent topological representation associated with every vertex *ni*. Φ is represented by an |N| × d matrix, where |N| is the cardinality of node set N, and d is the dimension of the node vector.

   $\Pr\left(n_i | \phi(n_i)\right)$ is approximated with *Hierarchial Softmax*[42] by assigning the nodes to the leaves of a Huffman tree, and $\Pr\left(n_i | \phi(n_i)\right)$ can be computed as:

$$\Pr\left(n_i | \phi(n_i)\right) = \prod_{l=1}^{\lceil \log |N| \rceil} \frac{1}{\left(1 + e^{-\phi(n_i)\psi(b_l)}\right)}$$

   where $b_l \in \left(b_0, b_{21}, .., b_{\lceil \log |N| \rceil}\right)$, and $\psi(b_l)$ is the representation assigned to the parent of node $b_l$. $\left(b_0, b_{21}, .., b_{\lceil \log |N| \rceil}\right)$ is a sequence of tree nodes to identify the node $n_i$, such that $b_0$ is the root of this tree and $b_{\lceil \log |N| \rceil} = n_i$.
3. *Computing embeddings:* After completing the above step, the latent topological representation of nodes in the network is the output of a hidden layer of the network.

DeepWalk performs its walks at random which means that embeddings do not preserve the local neighbourhood of the nodes well. However, other components of our pipeline, capture this local information from the virus and the mammalian perspectives.

By applying DeepWalk to compute the latent topological representation of our nodes, we can calculate the similarity between two nodes in our network n (vectorised as N) and m (vectorised as M) by using cosine similarity as follows [43,44]:

$$sim_{network}(n, m) = sim_{network}(M, N) = \frac{\sum_{i=1}^{d}(m_i \times n_i)}{\sqrt{\sum_{i=1}^{d} {m_i}^2} \times \sqrt{\sum_{i=1}^{d} {n_i}^2}}$$

where d is the dimension of the vectorised representation of our nodes: M and N, and $m_i$ and $n_i$ are the components of vectors M and N, respectively.

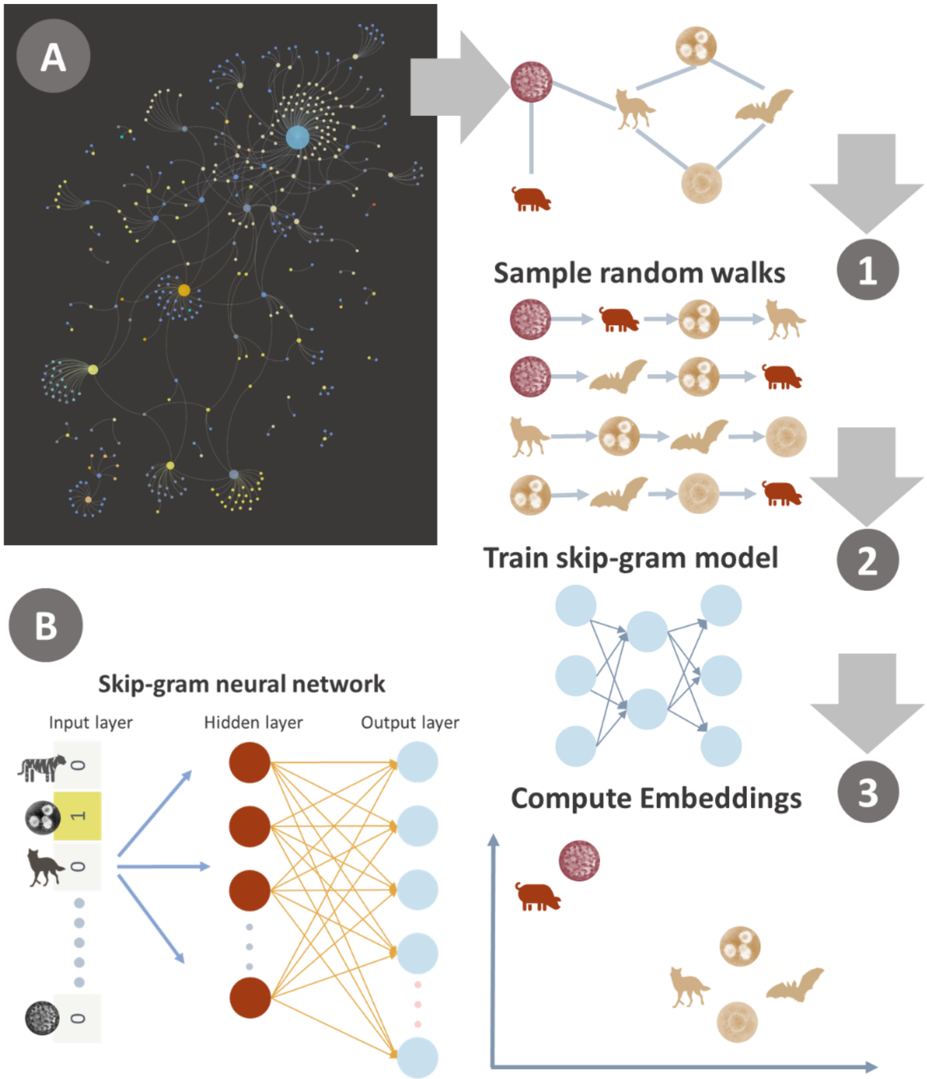


**Figure S3 – Network embeddings using DeepWalk. A – DeepWalk process. B Skip-gram neural network:** the network is composed of input layer, hidden layer(s) and output layer. The network accepts one-hot encoded vector. One-hot encoding is a vector with length same = number of nodes in network (or words in dictionary in the original implementation). The hidden layer has no activation function, its output presents an embedding of the node. The output layer is a softmax classifier that predicts neighborhoud nodes.

**Supplementary Note 4 – Similarity learning ensembles – multi-perspective approach**

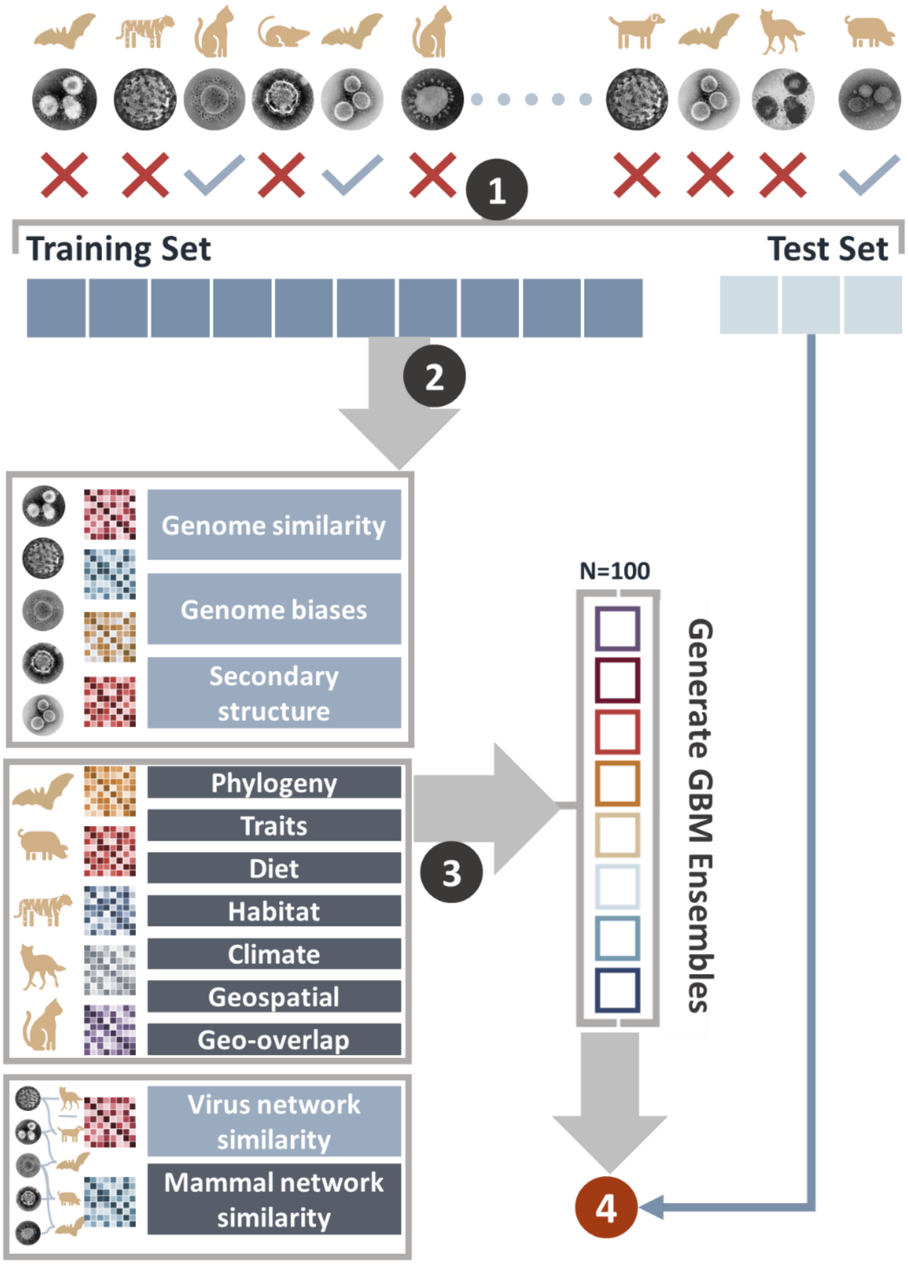


**Figure S4 – visualisation of training and performance assessment (5 repeats). Setp 1** – observed associations between CoVs and their hosts is split into training set comprising 85% of all observed associations – and test-set comprising 15% of these associations. **Setp 2** – training set is used to generate similarity learners in three categories: coronaviruses, mammalian hosts and networks. Network perspective learners are recomputed for each test run (from the reduced network). **Setp 3** – GBM is applied to generate meta-ensembles integrating the meta-learners. The ensembles comprises 100 replicate models trained with balanced samples drawn from the combined learners results. **Setp 4** – the model performance of the gbm ensemble is assessed by taking the mean probability of the 100 replicate models as applied to the test set.

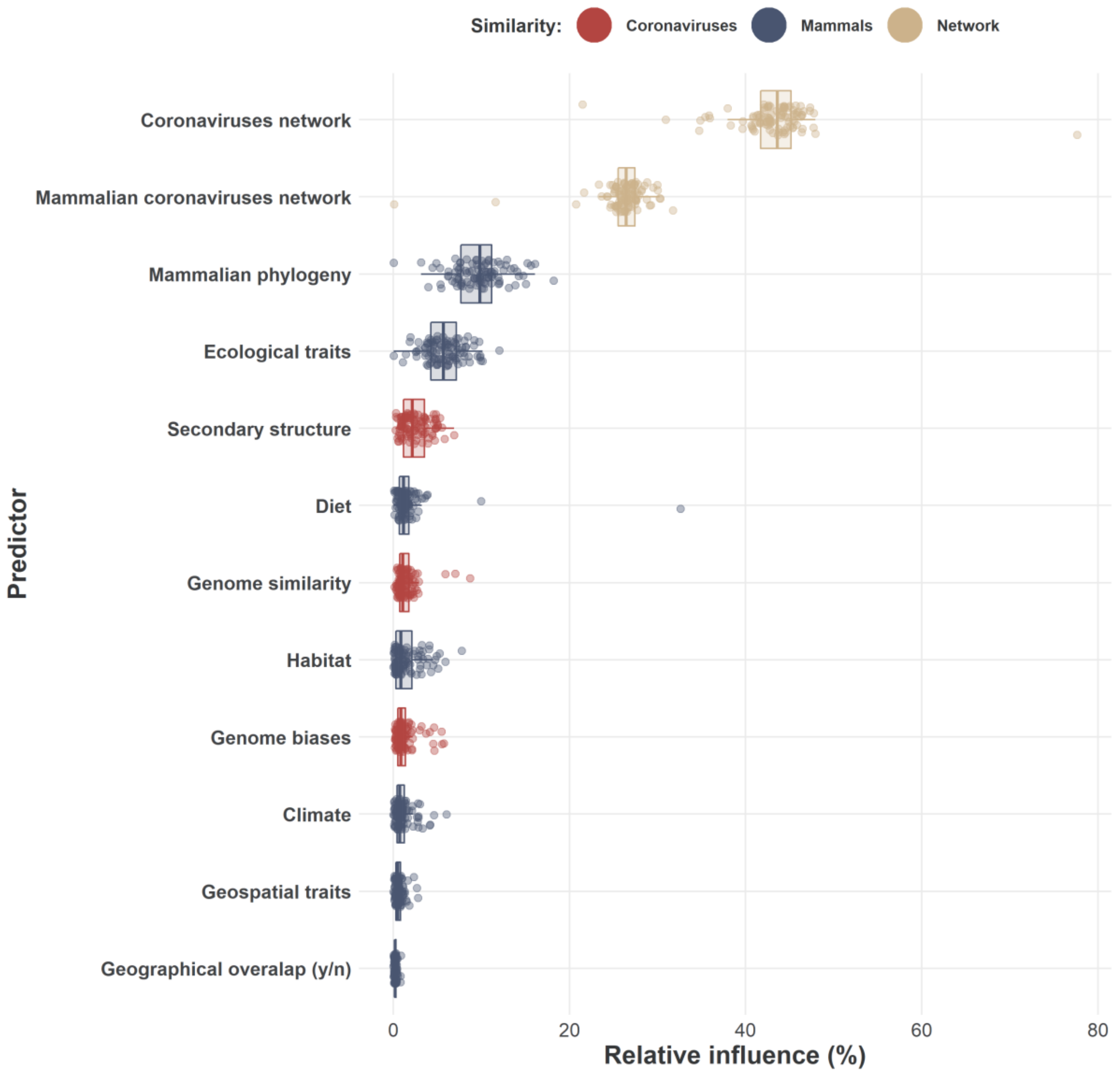


**Figure S5 – Relative influence (variable importance) of the included learners to the final gbm ensembles (trained with all available associations).**

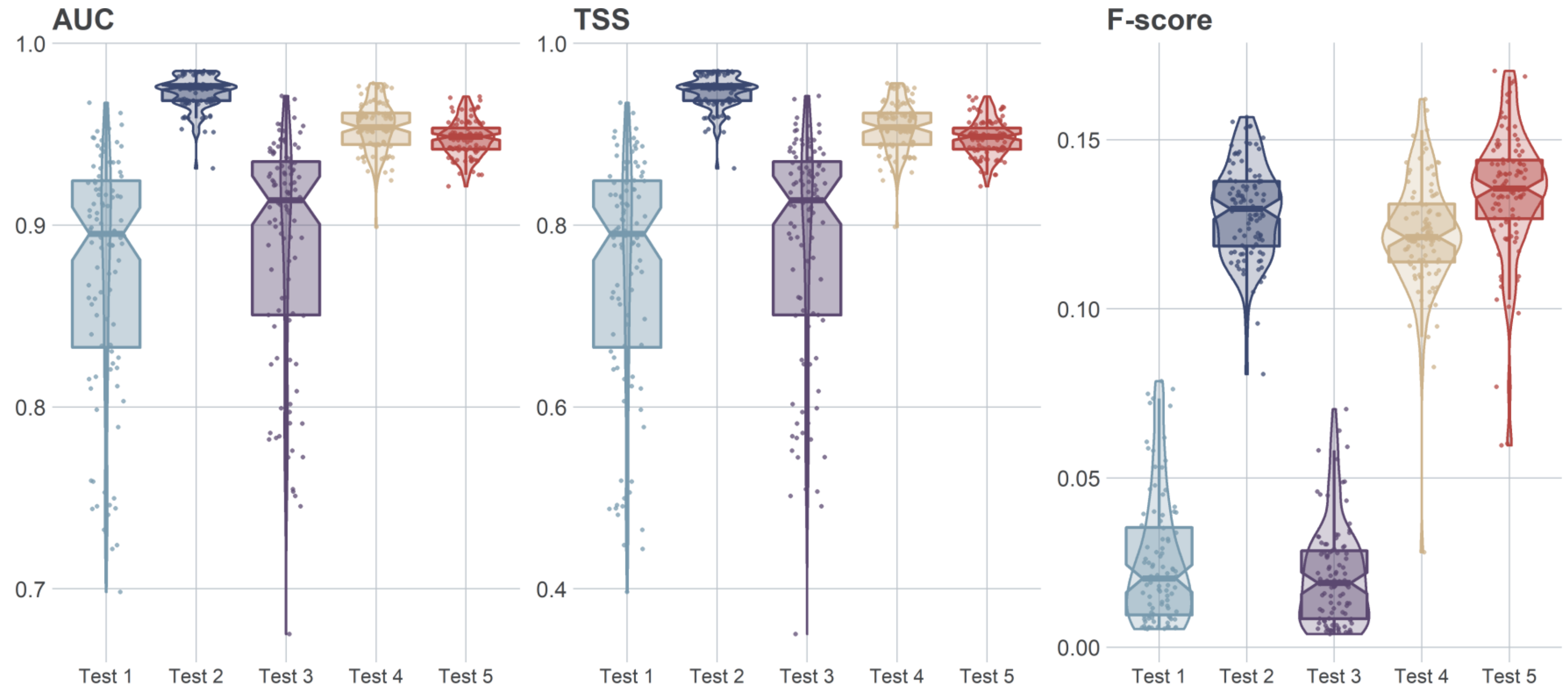


**Figure S6 – held-out test-set validation/performance assessment.**

#### Supplementary Note 5 – Changes in network structure with addition of predicted links

**Definitions**

| | |
|---|---|
| L | Number of realised links in the network. For our original network this number equals knowns associations between our CoVs and their mammalian hosts. In our predicted networks this number equals know and/or predicted links at the given probability cut-off. |
| M (J) | Number of mammalian species in the network. |
| V (I) | number of CoVs in the network |
| n (m) | total number of associations for all species |
| A | adjacency matrix of dimensions $\|V\| \times \|M\|$ where $\|V\|$ is number of coronaviruses included in this study (for which a complete genome could be found), and $\|M\|$ is number of included mammals, , such that for each $v_i \in V$ and $m_j \in M$, $a_{ij} = 1$ if an association exists (or is predicted at given cut-off) between the coronavirus and the mammal, and 0 otherwise. |
| $A_i$ | The total number of mammals associated with coronavirus $v_i \in V$. Such that, $A_i = \sum_{j=1}^{\|M\|} Aa_{ij}$. Corresponds to degree centrality (from the viral perspective). |
| $A_j$ | The total number of CoVs associated with mammal $m_j \in M$. Such that, $A_j = \sum_{i=1}^{\|V\|} Aa_{ij}$. Corresponds to degree centrality (from the viral perspective). |

Given the above definitions we computed the following structural properties at the level of the whole network, and the group (i.e. CoVs or mammals).

| Metric | Formula (method) | meaning |
|---|---|---|
| ***Mean degree*** | $\boldsymbol{D = \frac{L}{V+M}}$ | **Mean number of associations per CoV and mammal.** |
| *Connectance* | $C = \frac{L}{V \times M}$ | Realised proportion of possible associations (links). Deterministically increases with addition of new associations. |
| *Cluster coefficien*t | Mean per-node (CoV or mammal) connectance. Equals to mean, across all CoVs and mammals, of the number of realised associations (i.e. known and/or predicted) divided by the number of possible links for each node (CoV or mammal). | |
| *Cluster coefficien*t (CoVs) | Mean, across all CoVs, of the number of realised associations divided by the number of possible links for each CoV. | |
| *Cluster coefficien*t (mammals) | Mean, across all mammals, of the number of realised associations divided by the number of possible links for each mammal. | |
| Mean number of shared partners (CoVs) | Simple measure of co-occurrence. Capture mean number of shared hosts of CoVs. | |
| Mean number of shared partners (mammals) | Simple measure of co-occurrence. Capture mean number of shared CoVs between mammalian hosts. | |
| Togetherness (CoVs) | Togetherness *(CoVs)* measures the tendency of CoVs to be found in the same mammalian hosts. A high level (1) of togetherness (CoVs) suggests that the availability of a common trait or characteristics in these mammals (e.g. common receptor) might important in ability of CoVs to infect/association with them, whereas lower level (0) indicates the opposite[48,49]. | |
| Togetherness (mammals) | Togetherness *(mammals)* captures the tendency of mammalian species to shares CoVs. High values (max=1) of togetherness (mammals) suggest that similarities between mammalian species (e.g. habitat or diet requirements) might be more important driver of sharing of CoVs (community structure) than competition. Smaller values (min=0) indicate the opposite[48,49].. | |
| C-Score (CoVs) | Checkerboard score[50] (averaged | Measures non-independence in interaction patterns across the network. Larger values of C-Score (CoVs) indicates |

| | | |
|---|---|---|
| | across all nodes in level i.e. CoVs or mammals). | mammalian communities with little or no overlap in shared CoVs. |
| C-Score (mammals) | | Larger values of C-Score (mammals) suggest CoVs communities with little or no overlap in host preferences (e.g. tendencies of CoVs to be shared amongst certain host communities, defined, for example by phylogeny or geographical distribution). |
| V-ratio (CoVs) | variance-to-mean ratio | Larger values of this metric indicate a more skewed host range of our CoVs. |
| V-ratio (mammals) | | Larger values of this metric indicate a more skewed richness of CoVs in our mammalian hosts. |
| Nestedness | NODF *(nestedness metric based on overlap and decreasing fill)*[47] | Nestedness captures the tendency of specialists (e.g., CoVs with few hosts) to interact with (e.g. infects) subsets of mammals with which generalists (e.g., CoVs with many hosts) interact. It has been linked to network stability and functionality[45,46]. |
| Niche overlap (CoVs) | Mean similarity in interaction pattern between CoVs (in relation to mammalian species). Here we calculate this similarity via Horn's index (default implementation in the R package *bipartite*[51]). Niche overlap ranges from 0 (indicating no common pattern in how CoVs associate with mammalian species, to 1 indicating perfect overlap). | |
| Niche overlap (mammals) | Mean similarity in interaction pattern between mammals (in relation to CoVs). See above. | |
| Robustness (CoVs) | Area below the "*secondary extinction*" curve | CoVs are deleted at random, and area under the "second *extinction" curve is calculated*[51] *(see below).* |
| Robustness (mammals) | | Mammalian species are deleted at random, and area under the "second *extinction" curve is calculated*[51]. *Large values of robustness (max = 1) indicate* a curve that decreases very mildly until the point at which almost all mammalian species are eliminated. This suggests a very robust system in which, for instance, circulation of CoVs continues even if large fraction of mammalian host species is eliminated. On the other hand low values of robustness correspond to a curve that decreases abruptly as soon as any host species is lost. This is consistent with a fragile system in which, for instance, even if a very small fraction of the mammalian host is eliminated, most of CoVs lose their preferred hosts and drop from network (similar to extension events). |

**Table S2 –Network measures calculated for four bipartite networks (as presented in Figure 3): original network (3A), predicted network at probability cut off ≥0.95 (3B), >0.75 (3C), and >0.5 (3D) .** Values in bracket presents fold change. Calculations were based on adjacency matrices in which rows presented coronaviruses (higher level) and columns presented mammals (lower level). Cells values were 1 for associations (known or predicted at given cut-off) and 0 otherwise.

| **Metric** | original network | cut-off>=0.95 | cut-off>0.75 | cut-off>0.5 |
|---|---|---|---|---|
| **Mammalian diversity per virus** | 0.057 | 0.132 (0.087 - 0.515) / 2.316-fold (1.526 - 9.035) | 0.543 (0.131 - 0.548) / 9.526-fold (2.298 - 9.614) | 0.519 (0.331 - 0.595) / 9.11-fold (5.81 - 10.44) |
| **Viral diversity per mammal** | 0.241 | 0.666 (0.287 - 0.73) / 2.763-fold (1.191 - 3.029) | 0.731 (0.666 - 0.729) / 3.033-fold (2.763 - 3.025) | 0.73 (0.73 - 0.72) / 3.03-fold (3.03 - 2.99) |
| *Mean degree (associations per CoV or mammal)* | *1.055* | *3.449 (1.365 - 9.536) / 3.269-fold (1.294 - 9.039)* | *6.785 (3.433 - 11.41) / 6.431-fold (3.254 - 10.815)* | *9.398 (5.718 - 13.683) / 8.91-fold (5.42 - 12.97)* |
| *Connectance* | *0.011* | *0.028 (0.011 - 0.076) / 2.545-fold (1 - 6.909)* | *0.054 (0.027 - 0.091) / 4.909-fold (2.455 - 8.273)* | *0.075 (0.045 - 0.108) / 6.82-fold (4.09 - 9.82)* |
| *Cluster coefficient* | 0.005 | 0.023 (0.005 - 0.06) / 4.6-fold (1 - 12) | 0.041 (0.023 - 0.074) / 8.2-fold (4.6 - 14.8) | 0.06 (0.037 - 0.088) / 12-fold (7.4 - 17.6) |
| *Cluster coefficient (CoVs)* | 0.041 | 0.056 (0.036 - 0.127) / 1.366-fold (0.878 - 3.098) | 0.096 (0.056 - 0.153) / 2.341-fold (1.366 - 3.732) | 0.125 (0.08 - 0.182) / 3.05-fold (1.95 - 4.44) |
| *Cluster coefficient (mammals)* | 0.096 | 0.321 (0.166 - 0.532) / 3.344-fold (1.729 - 5.542) | 0.483 (0.318 - 0.539) / 5.031-fold (3.313 - 5.615) | 0.53 (0.447 - 0.549) / 5.52-fold (4.66 - 5.72) |
| **Nestedness (NODF)** | 6.065 | 38.758 (14.656 - 62.085) / 6.39-fold (2.416 - 10.237) | 56.856 (38.165 - 65.653) / 9.374-fold (6.293 - 10.825) | 61.705 (51.99 - 68.843) / 10.17-fold (8.57 - 11.35) |
| **Mean number of shared partners (CoVs)** | 0.207 | 1.905 (0.403 - 8.712) / 9.203-fold (1.947 - 42.087) | 5.627 (1.876 - 10.567) / 27.184-fold (9.063 - 51.048) | 8.555 (4.39 - 12.849) / 41.33-fold (21.21 - 62.07) |
| **Mean number of shared partners (mammals)** | 0.072 | 0.42 (0.097 - 2.798) / 5.833-fold (1.347 - 38.861) | 1.488 (0.419 - 4.056) / 20.667-fold (5.819 - 56.333) | 2.711 (1.026 - 5.855) / 37.65-fold (14.25 - 81.32) |
| **Cluster coefficient (CoVs)** | 0.041 | 0.056 (0.036 - 0.127) / 1.366-fold (0.878 - 3.098) | 0.096 (0.056 - 0.153) / 2.341-fold (1.366 - 3.732) | 0.125 (0.08 - 0.182) / 3.05-fold (1.95 - 4.44) |
| **Cluster coefficient (mammals)** | 0.096 | 0.321 (0.166 - 0.532) / 3.344-fold (1.729 - 5.542) | 0.483 (0.318 - 0.539) / 5.031-fold (3.313 - 5.615) | 0.53 (0.447 - 0.549) / 5.52-fold (4.66 - 5.72) |
| **Niche overlap (CoVs)** | 0.13 | 0.323 (0.229 - 0.567) / 2.485-fold (1.762 - 4.362) | 0.505 (0.318 - 0.579) / 3.885-fold (2.446 - 4.454) | 0.564 (0.459 - 0.589) / 4.34-fold (3.53 - 4.53) |

| | | | | |
|---|---|---|---|---|
| **Niche overlap (mammals)** | 0.053 | 0.065 (0.053 - 0.191) / 1.226-fold (1 - 3.604) | 0.149 (0.065 - 0.225) / 2.811-fold (1.226 - 4.245) | 0.187 (0.11 - 0.254) / 3.53-fold (2.08 - 4.79) |
| **Togetherness (CoVs)** | 0.007 | 0.043 (0.013 - 0.144) / 6.143-fold (1.857 - 20.571) | 0.094 (0.042 - 0.175) / 13.429-fold (6 - 25) | 0.14 (0.078 - 0.195) / 20-fold (11.14 - 27.86) |
| **Togetherness (mammals)** | 0.002 | 0.004 (0.001 - 0.018) / 2-fold (0.5 - 9) | 0.01 (0.004 - 0.026) / 5-fold (2 - 13) | 0.017 (0.007 - 0.036) / 8.5-fold (3.5 - 18) |
| **C-score (CoVs)** | 0.811 | 0.372 (0.65 - 0.122) / 0.459-fold (0.801 - 0.15) | 0.165 (0.381 - 0.117) / 0.203-fold (0.47 - 0.144) | 0.124 (0.213 - 0.104) / 0.15-fold (0.26 - 0.13) |
| **C-score (mammals)** | 0.931 | 0.81 (0.919 - 0.522) / 0.87-fold (0.987 - 0.561) | 0.61 (0.81 - 0.437) / 0.655-fold (0.87 - 0.469) | 0.531 (0.689 - 0.378) / 0.57-fold (0.74 - 0.41) |
| **V-ratio (CoVs)** | 7.207 | 9.057 (6.515 - 23.843) / 1.257-fold (0.904 - 3.308) | 17.781 (9.083 - 29.415) / 2.467-fold (1.26 - 4.081) | 23.29 (13.412 - 35.652) / 3.23-fold (1.86 - 4.95) |
| **V-ratio (mammals)** | 16.459 | 92.461 (43.033 - 156.718) / 5.618-fold (2.615 - 9.522) | 142.419 (90.995 - 158.811) / 8.653-fold (5.529 - 9.649) | 156.054 (131.029 - 163.227) / 9.48-fold (7.96 - 9.92) |
| **Robustness (CoVs)** | 0.538 | 0.58 (0.551 - 0.82) / 1.078-fold (1.024 - 1.524) | 0.724 (0.578 - 0.856) / 1.346-fold (1.074 - 1.591) | 0.818 (0.671 - 0.896) / 1.52-fold (1.25 - 1.67) |
| **Robustness (mammals)** | 0.586 | 0.791 (0.606 - 0.928) / 1.35-fold (1.034 - 1.584) | 0.896 (0.785 - 0.941) / 1.529-fold (1.34 - 1.606) | 0.926 (0.878 - 0.948) / 1.58-fold (1.5 - 1.62) |
